# Nemo knows: clownfishes differentiate cryptic host species across fine and broad geographic scales and reveal a potential adaptive radiation in the clownfish-hosting sea anemones

**DOI:** 10.1101/2024.11.15.623784

**Authors:** Tommaso Chiodo, Aurélien De Jode, Andrea Quattrini, Miranda K. Gibson, Catheline Y. M. Froehlich, Danwei Huang, Takuma Fujii, Kensuke Yanagi, James D. Reimer, Anna Scott, Estefanía Rodríguez, Benjamin M. Titus

## Abstract

The symbiosis between clownfishes (or anemonefishes) and their host sea anemones ranks among the most recognizable animal interactions on the planet. Found on coral reef habitats across the Indian and Pacific Oceans, 28 recognized species of clownfishes adaptively radiated from a common ancestor to live obligately with only 10 nominal species of host sea anemones. Are the host sea anemones truly less diverse than clownfishes? Did the symbiosis with clownfishes trigger a reciprocal adaptive radiation in sea anemones, or minimally, a co-evolutionary response to the mutualism? To address these questions, we combined fine- and broad-scale biogeographic sampling with multiple independent genomic datasets for the bubble-tip sea anemone, *Entacmaea quadricolor*—the most common clownfish host anemone throughout the Indo-West Pacific. Fine-scale sampling and restriction site associated DNA sequencing (RADseq) throughout the Japanese Archipelago revealed three highly divergent cryptic species: two of which co-occur throughout the Ryukyu Islands and can be differentiated by the clownfish species they host. Remarkably, broader biogeographic sampling and bait-capture sequencing reveals that this pattern is not simply the result of local ecological processes unique to Japan, but part of a deeper evolutionary signal where some species of *E. quadricolor* serve as host to the generalist clownfish species *Amphiprion clarkii* and others serve as host to the specialist clownfish *A. frenatus*. In total, we delimit at least five cryptic species in *E. quadricolor* that have diversified within the last five million years. The rapid diversification of *E. quadricolor* combined with functional ecological and phenotypic differentiation supports the hypothesis that this may represent an adaptive radiation in response to mutualism with clownfishes. Our data indicate that clownfishes are not merely settling in locally available hosts but recruiting to specialized host lineages with which they have co-evolved. These findings have important implications for understanding how the clownfish-sea anemone symbiosis has evolved and will shape future research agendas on this iconic model system.

## Introduction

The importance of mutualism is underscored by its ubiquity-virtually all of life engages in complex multi-level mutualisms that critically impact the formation and distribution of biodiversity around the globe (Margulis and Fester 1991; Doebeli and Knowlton 1998; Herre et al., 1999; Moran 2007; Bronstein 2015). The degree to which mutualistic partners exchange, or provide, resources should lead to variation in the degree to which selection mediates the symbiosis over evolutionary timescales (Herre et al., 1999; Bronstein 2015). Ultimately, these processes manifest themselves at various levels of biological organization, impacting the species, population, and genomic diversity of the constituent partners (reviewed by Bronstein 2015). Mutualisms can trigger adaptive species radiations, lead to rapid population expansions, and radically shape the architecture of the genome (Herre et al., 1999; Moran 2007; Moya et al., 2008; Mueller et al 2011; Brucker and Bordenstein 2012; Joy 2013; Bronstein 2015; Rubin and Moreau 2016). Yet the role of mutualism in generating biodiversity within symbiotic systems can be difficult to discern, largely because of the various evolutionary, geographic, and ecological scales in which they operate (Herre et al., 1999).

No mutualism is more representative of this puzzle than the iconic clownfish-sea anemone symbiosis, a model mutualism regularly used for exploring fundamental biological processes (Ollerton et al 2007; Litsios et al., 2012; Marcionetti et al., 2019; Sahm et al. 2019; Roux et al. 2020, 2024; Klann et al. 2021; Laudet and Ravasi 2022; Marcionetti and Salamin 2023 Moore et al. 2023), but one in which our understanding remains incomplete due to a lack of research into the evolution of the host sea anemones (De Jode et al., 2024; Titus et al., 2019a, 2024). Broadly distributed on Indo-Pacific coral reefs, 28 described species of clownfishes form obligate mutualisms with just 10 species of sea anemones (Anthozoa: Actiniaria) (Fautin and Allen 1992; Titus et al., 2024). Mutualism with sea anemones is considered the key ecological innovation that triggered the adaptive radiation of clownfishes ∼10-12 mya (Litsios et al., 2012), yet no evidence for host-driven patterns of diversification in clownfishes had been recovered in phylogenetic analyses until recently (De Jode et al., 2024; Gaboriau et al., 2024). Now, new evidence highlights that sea anemone host use can explain convergent phenotypic evolution in clownfish color patterns (Gaboriau et al., 2024), and further, that divergence times in the host sea anemones appear to be broadly coincident with the clownfish adaptive radiation (De Jode et al., 2024). The role of the host sea anemones in the evolution of the symbiosis is becoming increasingly established, yet key questions remain. Chief among these include: are the host sea anemones truly less diverse than the clownfishes? If not, given it is well established that the host anemones derive substantial benefits from hosting clownfishes, have the host sea anemones undergone their own hidden adaptive radiation?

One hypothesis is that the host sea anemones have undergone their own adaptive radiation, or are minimally, more diverse than currently recognized. Sea anemones have simple body plans, no hard parts, and few morphological characters that can reliably differentiate species (Titus et al. 2024). Undescribed cryptic species may thus be rampant within sea anemones (Titus et al. 2019a, b). Previous phylogenetic work using a suite of five traditional Sanger loci could not resolve species level relationships for the two most specious host clades Stichodactylina and Heteractina but were able to partially resolve two nodes within the clade *Entacmaea* Titus et al. 2019a). The population-level signal present in *Entacmaea* when using traditional mitochondrial and nuclear markers is atypical across Anthozoa (Shearer et al., 2002; Huang et al. 2008; Daly et al. 2010; McFadden et al. 2011; Quattrini et al. 2023), which hints that this nominal species may be a species complex.

*Entacmaea* Ehrenberg, 1834 is a monotypic genus in the sea anemone superfamily Actinioidea. The sole species, *E. quadricolor* (Leuckart in Rüppell & Leuckart, 1828), is colloquially referred to as the bubble-tip sea anemone due to the characteristic bubbles/bulges that form at the tentacle tips (Titus et al. 2024). Of the host sea anemones, *E. quadricolor* has one of the largest biogeographic ranges, spanning from the Northern Red Sea, throughout the Indian Ocean, Coral Triangle, and into the Pacific (Titus et al. 2024). It has been documented to serve as a host to 20 of the 28 clownfish species, including both generalist and specialist clownfish species (Gaboriau et al., 2024). Where multiple clownfish species co-occur on the same reefs, *E. quadricolor* is regularly observed as hosts for multiple species (Fautin and Allen 1992).

*Entacmaea quadricolor* also displays a great deal of intraspecific morphological variation in color, pattern, and tentacle shape (Titus et al. 2024). This has historically been attributed to phenotypic plasticity rather than species level differences (Fautin and Allen 1992; Richardson et al 1997; Titus et al. 2024). The degree of ecological and morphological variation within this single nominal species makes *E. quadricolor* an ideal candidate to test the degree to which the clownfish-hosting sea anemones are under described. The global importance of *E. quadricolor* as a host to >70% of all clownfish species further makes this putative species ideal for testing whether there are signatures of adaptive radiation within the host anemones.

Here we combine fine- and broad-scale biogeographic sampling of *E. quadricolor* with two independently derived genomic datasets to test for undescribed cryptic lineages and potential signatures of adaptive radiation linked to their mutualism with clownfishes. We couple our genomic datasets with an assessment of phenotypic diversity to tease apart any potential morphological differences among cryptic lineages. We discover that *E. quadricolor* is a diverse species complex that has rapidly diversified within the last five million years. Remarkably, we find co-occurring cryptic lineages of *E. quadricolor* not only host different species of clownfishes on the same reefs in the Japanese Archipelago, but are phenotypically different and derived from deeper evolutionary lineages linked to hosting specialist and generalist clownfish species. Our findings reveal far greater specialization and patterns of co-evolution in the clownfish-sea anemone symbiosis and the first potential evidence of a reciprocal adaptive radiation in the hosts. These findings will radically alter future research agendas within this iconic mutualism.

## Materials and Methods

### Sample collection

To search for patterns of cryptic species-level diversity within *Entacmaea quadricolor*, we first conducted fine-scale phylogeographic surveys and sampling across the Japanese Archipelago. We surveyed and photographed N = 126 individuals and collected N = 93 tentacle clippings from 38 sample localities, including the remote Ogasawara Islands ∼1000 km from Mainland Japan (Figure 1; Table S1). Samples were collected by hand using SCUBA between 1-29 m depth. *In situ* photos were taken of each anemone, clownfish symbionts were identified and quantified, and one or two tentacles were sampled using forceps and placed into individually labeled collection bags. On shore, samples were preserved in 95% ethanol. To place our Japanese samples into a broader biogeographic context, we also collected *E. quadricolor* samples from adjacent biogeographic regions including Singapore, Maldives, Australia, and the Philippines (Table S3). Samples were collected and preserved as above.

**Figure 1.**
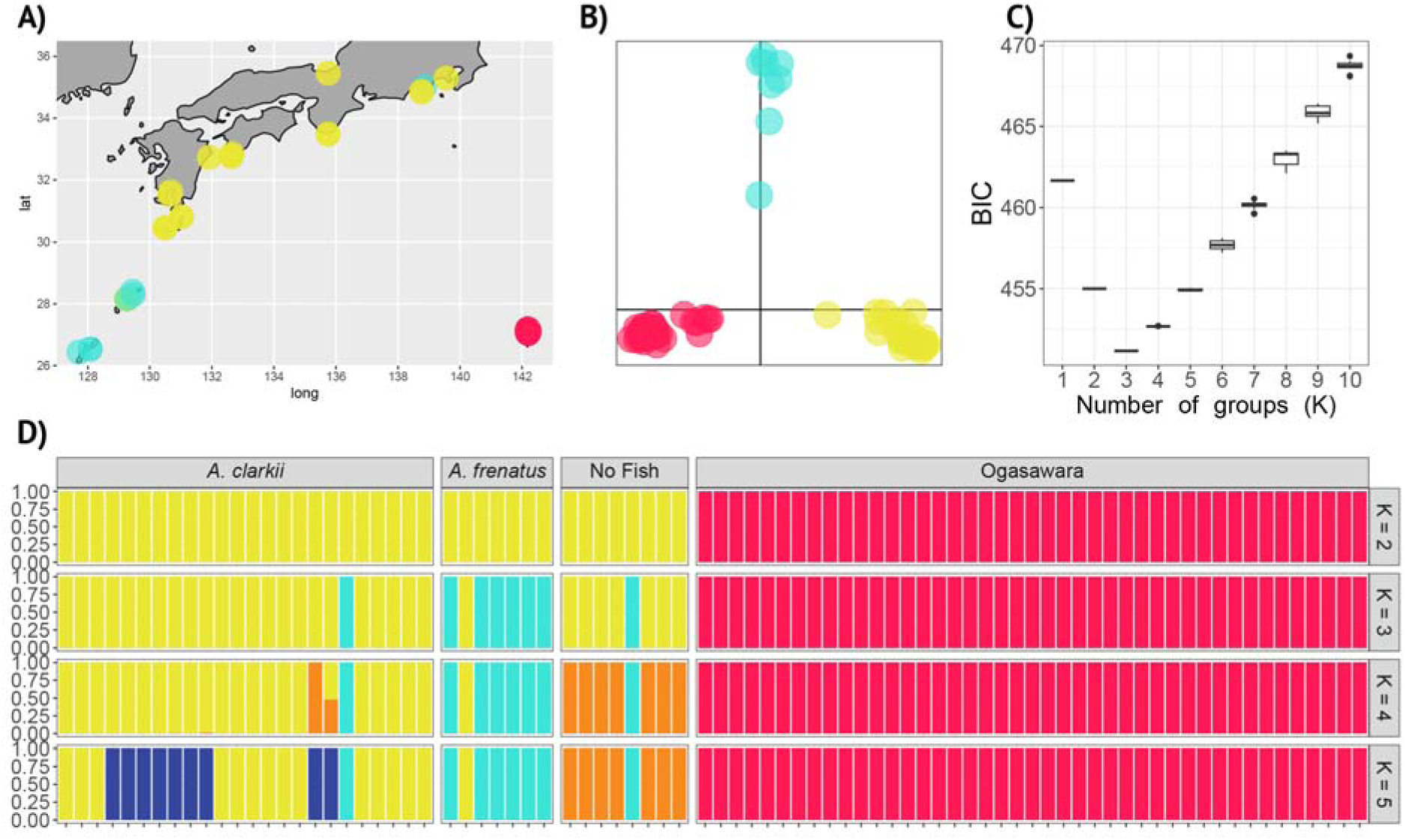
Species delimitation results for the bubble-tip sea anemone *Entacmaea quadricolor* using double-digest restriction site associated DNA sequencing (ddRADseq) demonstrating cryptic lineages of *E. quadricolor* co-occur throughout Japan and host different species of clownfishes. A) Field sampling localities across the Japanese Archipelago and the Ogasawara Islands (Red). Sampling localities are color coded by the results of discriminant analysis of principal components (DAPC) genetic clustering results for each individual *E. quadricolor* sea anemone in the dataset (panels B and D) and reflect newly delimited cryptic *E. quadricolor* species. B) DAPC scatter plot of axis one and two for *k* = 3 genetic cluster assignments. C) Results of Bayesian Information Criterion model selection results demonstrating *k* = 3 as the best genetic clustering model for *E. quadricolor*. D) Posterior probabilities of individual group assignments for *k* = 2-5 for the ddRADseq dataset. For co-occurring cryptic *E. quadricolor* lineages in the Ryukyu Islands and Mainland Japan (yellow and teal) the identity of the clownfish symbiont present in each anemone is provided.

### DNA Extraction, Library Preparation, and Sequencing

Following collection, total genomic DNA was extracted using a modified DNeasy Blood and Tissue Kits (QIAGEN Inc.) and prepared for DNA sequencing (see Supplementary Methods). To test for undescribed cryptic diversity within *E. quadricolor* and place the resulting diversity into broader phylogeographic context, we generated two independent genomic datasets. First, for all samples we collected throughout the Japanese Archipelago, we used a double-digest restriction-site associated DNA sequencing (ddRADseq) approach for cryptic species discovery (Titus et al. 2019b; Titus et al., 2022; see Supplementary Methods). Next, we used bait-capture sequencing targeting ultra-conserved elements (UCEs) and exon loci to produce a second genomic dataset and place the resulting cryptic lineages recovered from Japan into hierarchical biogeographic context. Accordingly, we tested whether the *E. quadricolor* lineages in Japan were diversifying *in situ* or were the result of geographical range overlap of more distantly related species. We re-sequenced N = 64 individuals from Japan along with N = 46 additional samples from adjacent biogeographic regions including Singapore (Chan et al. 2020), Philippines, Australia, and Maldives (Table S2). We used this alternative genomic approach as *E. quadricolor* does not have a closely related sister species that would be suitable to use as an outgroup taxon to root a ddRADseq dataset (Titus et al. 2019a; De Jode et al. 2024) and previously published bait-capture data for sea anemones are publicly available on GenBank (Table S3) (Quattrini et al. 2018, 2020). Bait-capture libraries were prepared and sequenced following the protocol developed by Quattrini et al. (2018; see Supplementary Methods).

### Dataset assembly

Following sequencing, raw ddRADseq data were demultiplexed, aligned, and assembled *de novo* using ipyrad (Eaton and Overcast 2020). We set the clustering threshold (Wclust) to 0.90 to assemble reads into loci and sequencing depth (e.g. mindepth_statistical and mindepth_majrule) parameters were set to 10 to reduce heterozygous calls due to sequencing error. The min_samples_locus was set to 76 as that would represent approximately 75% occupancy across samples for the locus to be retained in the ddRADseq dataset. One SNP per locus was randomly selected to build an unlinked SNP dataset. Raw ddRADseq data were deposited at the NCBI Sequence Read Archive under BioProject XXXXX (Table S1).

To assemble UCE and exon loci from raw bait-capture sequence data we used the program PHYLUCE (Faircloth 2016) and the hexa_v2_final bait set (Cowman et al. 2020; Glon et al. 2020). After demultiplexing, raw sequences for each individual sample were cleaned and trimmed using illumiprocessor (Bolger et al. 2014; Faircloth 2016, 71). Cleaned sequences were then assembled into contigs using SPAdes v3.14.1 (Bankevich et al. 2012) with the -careful and - cov-cutoff 2 parameters. We then used PHYLUCE, as described in online tutorials to extract exon and UCE loci and assemble the final dataset. UCE baits were matched and then extracted using phyluce_assembly_ match_contigs_probes and phyluce_assembly_get_match_counts. We created a final dataset with 75% completeness (i.e. no more than 25% missing data at a single locus). Our dataset was aligned using MAFFT (Katoh et al. 2002) and internally trimmed using the version of Gblocks inside PHYLUCE (phyluce_align_get_gblocks_trimmed_alignments_from_untrimmed) with the default parameters. For downstream bait-capture analyses, *Epiactis georgiana* Carlgren, 1927 was used as an outgroup species as this is currently the most closely related known species to *E. quadricolor* (De Jode et al. 2024).

### Fine scale phylogeography and species delimitation in the Japanese Archipelago

To test whether there is evidence that *E. quadricolor* is a cryptic species complex within the Japanese Archipelago we used our fine-scale ddRADseq dataset to conduct three species discovery analyses. We first conducted a principal coordinates analysis (PCA) as a preliminary investigation to visualize the genetic variation present in the Japanese Archipelago in our unlinked SNP dataset generated by ipyrad (see Supplementary Methods). We next performed a Discriminant Analysis of Principal Components (DAPC) on the ddRADseq data to assign individuals of *E. quadricolor* from Japan to genetic clusters using the R package adegenet (Jombart 2008). The optimal value of *k* was obtained using a *k*-means clustering algorithm (find.clusters()) and the *k* value with the lowest BIC value was selected. To determine the number of principal components to retain for the DAPC, an 12-score optimization run was done and cross validated by selecting the number of PCs with the highest average prediction success per group. A separate DAPC analysis was conducted for *k* = 2-5 to compare group assignments of different *k* values to our biological lineages. Finally, we used the Bayesian genetic clustering approach implemented in the program fastStructure (Raj et al. 2014) to delimit major genetic clusters and assign individuals to putative species. Using default parameters, we tested *k* values from 2 to 5 to determine number of genetic clusters that maximized the marginal likelihood for our dataset. All clustering programs identified the same three major genetic clusters, and we computed pairwise F_ST_ to determine the degree of genetic differentiation within and between clusters using the R package hierfstat (Goudet 2005).

### Entacmaea quadricolor phenotypic diversity

To determine whether our newly delimited species in the Japanese Archipelago displayed consistent phenotypic characteristics that could be used to visually identify each lineage apart from hosting different clownfishes, we phenotyped N = 126 *E. quadricolor* individuals from field survey photographs. For each individual, we recorded phenotypic information for tentacle shape, tentacle length, tentacle tip pattern, tentacle color, tentacle tip color, anemone group size, depth, and finally, clownfish symbiont species. Tentacle shape variables were categorized as “rounded bubble tips,” “bubble with extended tip,” or digitiform (uniformly shaped/finger-like tentacles with no bubble tip swelling; Figure 2). Tentacle length was categorized as short (< 5 cm) or long (> 5 cm). Tentacle tip pattern was categorized into Dull/Matte (D/M), striated (ST), or striated and speckled (STSP; Figure 2). Tentacle color was categorized into bleached, brown, pale brown, green/brown, and green. Tentacle tip color was categorized as whether the tentacle tips had pink tips (PT) or not (noPT). Anemone group size was categorically defined as whether the anemone was a solitary individual (S) or part of a clonal group (G). To account for important differences in habitat, we included depth as a variable. Depth was binned into shallow (< 5m) or deep (>5m). We recorded whether the clownfish present in the anemone was *A. clarkii*, *A. frenatus*, or if no fish were present. We tested for significant associations between each phenotypic character and our three newly delimited species using multinomial logistic regression models. Analyses were conducted with the vglm function in R-Studio using packages tidyverse (Wickham et al. 2019), VGAM (Yee 2010), car (Fox and Weisberg 2018), and rcompanion (Mangiafico 2016).

**Figure 2.**
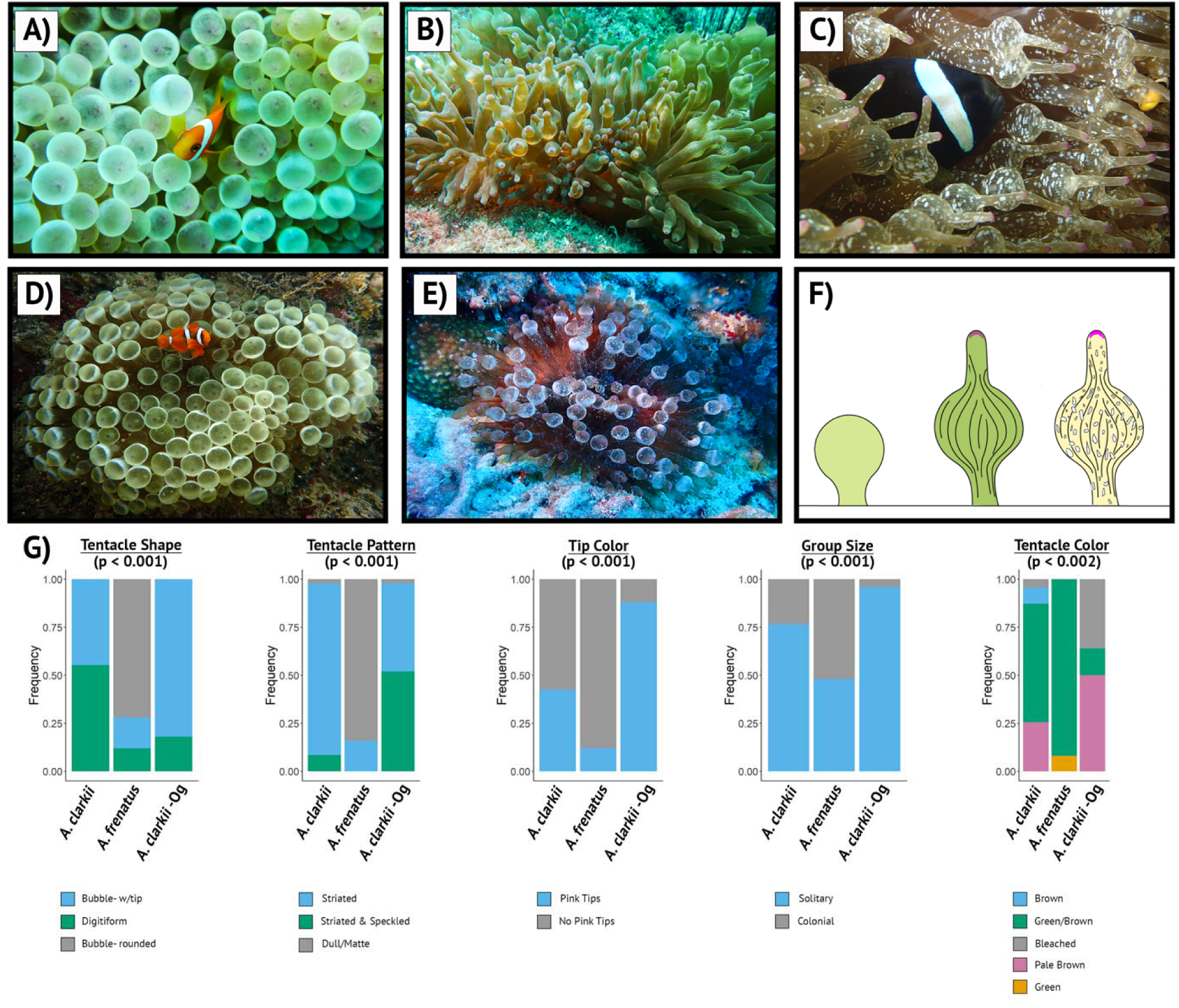
Consistent phenotypic differences among cryptic *Entacmaea quadricolor* species that host different clownfishes in the Japanese Archipelago. A) *Amphiprion frenatus* hosting sea anemones from the Ryukyu Islands are characterized by rounded bubble tips, a dull/matte tentacle tip pattern, and no significant terminal tentacle tip color. B) *A. clarkii* hosting sea anemones from the Ryukyu Islands and Mainland Japan have bubble-tip tentacles with an extended tip protruding from the bulb and a striated tentacle tip pattern. C) *A. clarkii* hosting sea anemones from the Ogasawara Islands have bubble-tip tentacles with an extended tip protruding from the bulb, a striated and speckled tentacle tip pattern, and are frequently a pale brown color. D) Representative photograph (by Y.C. Tay) of a bubble-tip sea anemone from Singapore that hosts *A. frenatus* and has a qualitatively similar phenotype to the *A. frenatus* hosting anemones from Japan. E) Representative photograph of a bubble-tip sea anemone from the Maldives that hosts *A. clarkii* and has a qualitatively similar phenotype to the *A. clarkii* hosting anemones from Mainland Japan. F) Representative tentacle shape, color, and pattern that align with the three cryptic *E. quadricolor* species in panels A, B, and C, respectively.

### Phylogenetic reconstruction

To place newly delimited *E. quadricolor* species from Japan into hierarchical and biogeographic context, we conducted phylogenetic analyses using our bait-capture sequencing dataset targeting UCE and exon loci. Using our 75% dataset, we constructed Maximum Likelihood (ML) phylogenetic analyses using IQTree2 (Minh et al 2020). IQTree2 analyses were conducted using a concatenated and partitioned dataset with nodal support assessed using 10,000 ultrafast bootstraps (Hoang et al. 2018) and SH-like approximate likelihood ratio tests (Guindon et al. 2010). Partitioned IQTree2 analyses were generated using the loci positions output by PHYLUCE to delimit partitions and ModelFinder to select the most appropriate substitution model (Chernomer et al. 2016; Kalyaanamoorthy et al. 2017).

Next, we used a coalescent-based approach to reconstruct phylogenetic relationships in ASTRAL III (Zhang et al. 2018). ASTRAL III accounts for incomplete lineage sorting by estimating a species tree from many independent gene trees. Input gene trees were generated using IQTree2 for each locus in the 75% occupancy matrix using the best fit model of evolution selected with ModelFinder (Kalyaanamoorthy et al. 2017). The resulting trees were used as an input for ASTRAL-hybrid v1.15.2.3 analyses. Nodal support in ASTRAL III analyses were assessed using posterior probabilities. Lastly, we reconstructed *E. quadricolor* relationships using CASTER (Zhang et al. 2023). CASTER represents another coalescent-based species-tree approach but uses a multiple sequence alignment as input instead of gene trees. CASTER analyses were run with the concatenated 75% multi-sequence alignment and nodal support evaluated using posterior probabilities.

To estimate divergence times for newly delimited *E. quadricolor* species recovered from our phylogenomic analysis, we constructed a large UCE dataset by combining our *E. quadricolor* UCE dataset with previously published bait-capture sequence data from Order Actiniaria (Table S3), including samples from all major sea anemone superfamilies. We conducted a full analysis of all Actiniaria to capture the root of all sea anemones as the root age for Actiniaria had been previously estimated in a broader fossil calibrated phylogenomic analysis of Class Anthozoa (Quattrini et al. 2020; McFadden et al. 2021) A time calibrated dataset was assembled following the same modified PHYLUCE pipeline to generate a 75% occupancy matrix and IQTree2 was used to generate an unpartitioned ML tree as the unpartitioned tree retained higher support values for deeper relationships. Using IQTree2, divergence times were estimated by converting substitutions per site in the unpartitioned ML tree to years by setting the root age of Actiniaria to the previously estimated divergence date between 424 and 608 million years ago (Ma) (Quattrini et al. 2020; McFadden et al 2021). The tip date was set to 0 and the confidence interval to 1,000 years.

### Demographic modeling

We conducted demographic modeling in *dadi* (diffusion approximations for demographic inference; Gutenkunst et al. 2009) to test competing diversification scenarios involving our newly delimited *E. quadricolor* from the Japanese Archipelago (*A. frenatus*-hosting, *A. clarkii*-hosting, and Ogasawara Islands). We tested whether ancestral diversification between *A. frenatus* and *A. clarkii* lineages occurred with or without geneflow, as well as tested whether co-occurring lineages in Japan were reproductively isolated. We built a set of 17 demographic models (Figure 4) following protocols and pipelines detailed by Portik et al. (2017). Each model was a three-population isolation-migration model that varied in the directionality and timing of gene flow between *E. quadricolor* species.

To model the evolutionary history of *E. quadricolor*, we built a three-dimensional Joint-folded Site Frequency Spectrum (JSFS) using our ddRADseq dataset from the Japanese Archipelago. Individual anemones were assigned to putative species based on species delimitation results. All loci not in Hardy-Weinberg Equilibrium (p > 0.05) were removed using VCFTools (Danecek et al. 2011). Model simulations were conducted with consecutive rounds of optimization, where multiple replicates and previous parameter estimates from best scoring replicates were used to seed subsequent simulations. The default dadi_pipeline settings were used for each round (replicates = 10, 20, 30, 40; maxiter = 3, 5, 10, 15; fold = 3, 2, 2, 1) and the parameter optimization followed the Nelder-Mead method (optimize_log_fmin). The optimized parameters of each replicate were used to simulate the 3D-JSFS and estimate the log-likelihood of the JSFS of the model. The best fit demographic model was selected using Akaike Information Criterion and model probabilities were calculated following Burnham and Anderson (2002).

## Results

### Fine scale phylogeography and species delimitation in the Japanese Archipelago

Fine-scale sampling and ddRADseq sequencing efforts resulted in 202 million raw sequence reads. After assembly and filtering in *ipyrad* our final dataset contained 3,516 unlinked loci across 82 individuals. All species discovery methods consistently recovered three distinct *E. quadricolor* lineages (Figure 1). Principal Components Analysis (PCA), *k*-means clustering using discriminant analysis of principal components analysis (DAPC), and fastStructure all recovered two co-occurring *E. quadricolor* lineages in the Ryukyu Islands and a third allopatric lineage from the Ogasawara Islands (Figure 1; Figure S1 & S2).

Surprisingly, where the two putative *E. quadricolor* lineages co-occurred throughout the Ryukyu Islands they hosted different species of clownfishes with near 100% perfect habitat segregation. Our most heavily sampled *E. quadricolor* lineage harbored the host generalist Clark’s clownfish *Amphiprion clarkii*. This species occurred from the Ryukyu Islands in southern Japan north into Mainland Japan. Our less frequently sampled *E. quadricolor* lineage harbored the host specialist Tomato clownfish *A. frenatus,* which is known to exclusively associate with *E. quadricolor* (Figure 1; Figure S1 & S2). We only sampled the *A. frenatus* hosting lineage in the Ryukyu Islands. In our most northern sampling localities from Mainland Japan, we collected some *E. quadricolor* individuals from shallow habitats (<1m depth) that did not host clownfishes. These clustered with the putative *A. clarkii* hosting lineage (Figure 1). In Ogasawara, our third lineage also served as host to *A. clarkii* although the Ogasawaran clownfishes exhibit a unique all-black phenotype and there has been speculation these represent a cryptic species within the *A. clarkii* complex. Genetic divergence between putative *E. quadricolor* lineages was high (Table 1). Pairwise Fst values were higher between co-occurring *E. quadricolor* lineages that hosted different clownfishes than between *E. quadricolor* lineages that hosted *A. clarkii* clownfishes but where allopatrically distributed between Mainland Japan and Ogasawara Islands (Table 1).

**Table 1.** Pairwise F_ST_ values between newly delimited cryptic species of *Entacmaea quadricolor* (EQ) in the Japanese Archipelago that host different clownfishes (*Amphiprion frenatus* and *A. clarkii*). F_ST_ values were calculated using double-digest restriction site associated DNA sequencing (ddRADseq).

|  | EQ-*A. frenatus* | EQ-*A. clarkii* |
| --- | --- | --- |
| EQ-*A. frenatus* | - |  |
| EQ-*A. clarkii* | 0.27 | - |
| EQ-Ogasawara | 0.32 | 0.15 |

### Entacmaea quadricolor *phenotypic diversity*

Phenotypic analyses revealed that tentacle shape, tentacle tip pattern, tentacle tip color, anemone group size, and tentacle color were all predictive of the three cryptic host species (Figure 2). In general, *E. quadricolor* anemones that hosted the Tomato clownfish *A. frenatus* had bubble-tip tentacle shapes with blunt-rounded tips and a dull-matte tentacle tip pattern (Figure 2a, f). These could be differentiated from co-occurring *E. quadricolor* anemones that hosted *A. clarkii,* which had bubble-tip tentacle shapes with a protruding/elongated tentacle tip extending from the bulbous swelling and had striated (i.e. striped) tentacle tip patterns with pink tips (Figure 2b, f). Ogasawaran *E. quadricolor* anemones had similar phenotypes as the *A. clarkii* hosting anemones in Mainland Japan with the addition of regularly having a speckled tentacle tip pattern and a pale-brown coloration (Figure 2c, f). The *A. frenatus* hosting *E. quadricolor* were also significantly more likely to be found in groups/aggregations of anemones and were less likely to have pink tipped tentacles, in contrast to both *A. clarkii* and Ogasawaran anemones, which were more frequently found solitarily and regularly had pink tentacle tips (Figure 2).

### Phylogenetic Reconstruction

Bait-capture sequencing and dataset assembly resulted in an average of 56,630±48,647 contigs per sample with a mean locus length of 329±85 basepairs across N = 103 individuals from Japan, Singapore, Australia, Maldives, and the Philippines (Table S3). The total average base pairs per sample was 18,603,577±12,788,845. After alignment and edge trimming, we recovered 1,542 UCE and exon loci. We assembled a final dataset requiring at least 75% completeness per locus, resulting in 1002 retained loci and 88,410 parsimoniously informative sites.

Partitioned maximum likelihood phylogenetic analyses in IQtree2 resulted in a highly supported tree topology (Figure 3). We recovered the *E. quadricolor* lineage from the Ryukyu Islands and Mainland Japan that hosted *A. clarkii* as sister to the *A. clarkii* hosting *E. quadricolor* lineage from Ogasawara (Figure 3). Interestingly, we recovered *E. quadricolor* individuals from the Maldives, which also hosted *A. clarkii*, as sister to both Japanese *E. quadricolor* lineages that hosted *A. clarkii.* Additionally, we recovered the *E. quadricolor* lineage from the Ryukyu Islands that hosts *A. frenatus* to form a sister relationship with *E. quadricolor* from Singapore, which also hosts *A. frenatus*. Thus, the *E. quadricolor* species complex in our analyses formed two deep evolutionary clades associated with hosting generalist (*A. clarkii*) and specialist (*A. frenatus*) clownfishes (Figure 3). IQtree2 phylogenetic analyses placed all samples from Australia in a deeper monophyletic clade sister to all other samples (Figure 3). Individual *E. quadricolor* anemones from the Solitary Islands (Australia) hosted the Great Barrier Reef clownfish *A. akindynos*. The clownfish identity from the *E. quadricolor* anemones collected from the Great Barrier Reef were unknown as these were provided by Cairns Marine Inc. in the aquarium trade. In total, IQtree2 phylogenetic analyses recovered seven highly supported monophyletic *E. quadricolor* lineages that correspond to either clownfish host use or geography: 1) Ryukyu Islands and Mainland Japan – *A. clarkii,* 2) Ogasawara Islands – *A. clarkii*, 3) Maldives – *A. clarkii*, 4) Ryukyu Islands Japan – *A. frenatus*, 5) Singapore – *A. frenatus*, 6) Great Barrier Reef Australia, 7) Solitary Islands Australia (Figure 3).

**Figure 3.**
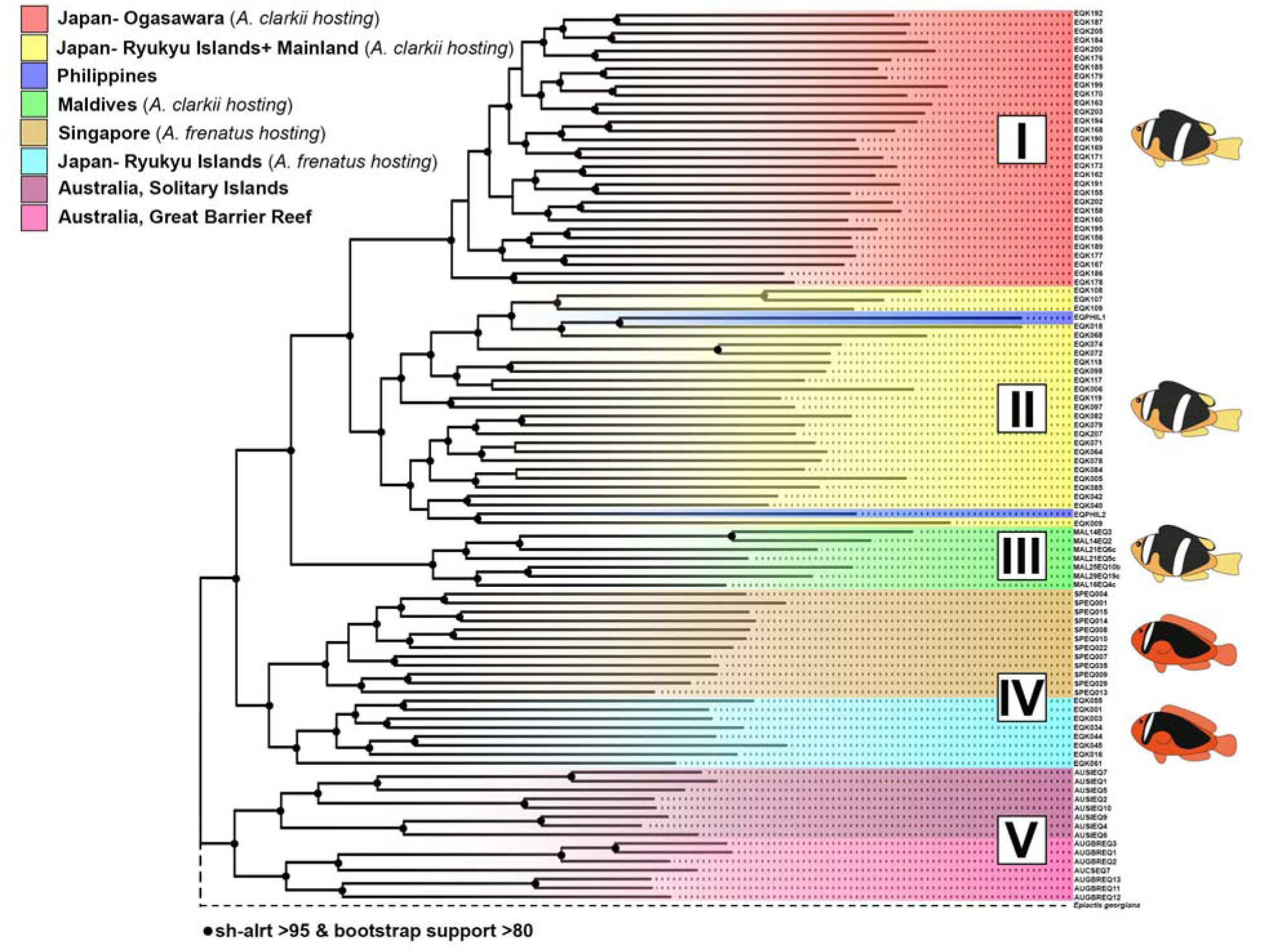
Maximum Likelihood (ML) phylogenetic reconstruction of the bubble-tip sea anemone *Entacmaea quadricolor* reveals at least five cryptic species (I-V) and that generalist and specialist clownfishes segregate by cryptic host sea anemone species over broad geographic scales. ML analyses conducted using partitioned phylogenetic analysis in IQtree2 and bait-capture dataset targeting ultra-conserved element and exon loci (75% data occupancy matrix and 1002 loci). Results demonstrate that co-occurring *E. quadricolor* lineages in Japan are not the result of *in situ* diversification in Japan, but rather geographic range overlap of more deeply divergent lineages linked to the clownfish species they host.

Coalescent-based phylogenetic analyses in ASTRAL-III and CASTER recovered similar *E. quadricolor* lineages and tree topologies as IQtree2 but with some differences (Figure S3 & S4). Importantly, all analyses recovered the same three cryptic Japanese *E. quadricolor* lineages and demonstrated that co-occurring Japanese diversity was not the result of *in situ* endemic diversification (Figure S3 & S4). Some ambiguity existed in the placement of the *E. quadricolor* lineage from the Maldives. In ASTRAL-III, the Maldives were placed as sister to Australia, Singapore, and the *A. frenatus* hosting *E. quadricolor* from Japan, rather than to the *A. clarkii* hosting anemones from Japan and the Ogasawaran Islands (Figure S3). Deep nodes, however, were less well supported in ASTRAL-III analyses than from IQtree2. CASTER phylogenetic analyses placed the Maldives back as sister to both *A. clarkii* hosting lineages from Japan with full support (Figure S4).

To estimate divergence times for newly delimited *E. quadricolor* species, we incorporated an additional N = 84 previously sequenced sea anemone samples to capture the root of Order Actiniaria. We recovered 2496 UCE and exon loci with 185,085 parsimoniously informative sites. Using the previously estimated 95% confidence intervals for the root age for Order Actiniaria (424-608 Ma) our time calibrated analysis in IQtree2 dated divergence times for all members of the *E. quadricolor* species complex to be between 5.7 and 3 Ma (Figure S5).

### Demographic Modeling

Among 17 unique demographic models, Akaike Information Criterion (AIC) model selection placed all model support on a three-population isolation-migration model (Figure 4, Table S3). The model included divergence with ancestral, bi-directional, symmetrical, migration at time 1 (T1), and divergence with contemporary unidirectional migration from Ryukyu Islands/Mainland *A. clarkii* hosting *E. quadricolor* to Ogasawaran *E. quadricolor* at T2 (Figure 4; Table S3). Our best fit model did not include a gene flow parameter between co-occurring *E. quadricolor* anemones that host different clownfishes on reefs in the Ryukyu Islands during T2. No other simulated demographic model garnered appreciable model support (Table S3).

**Figure 4.**
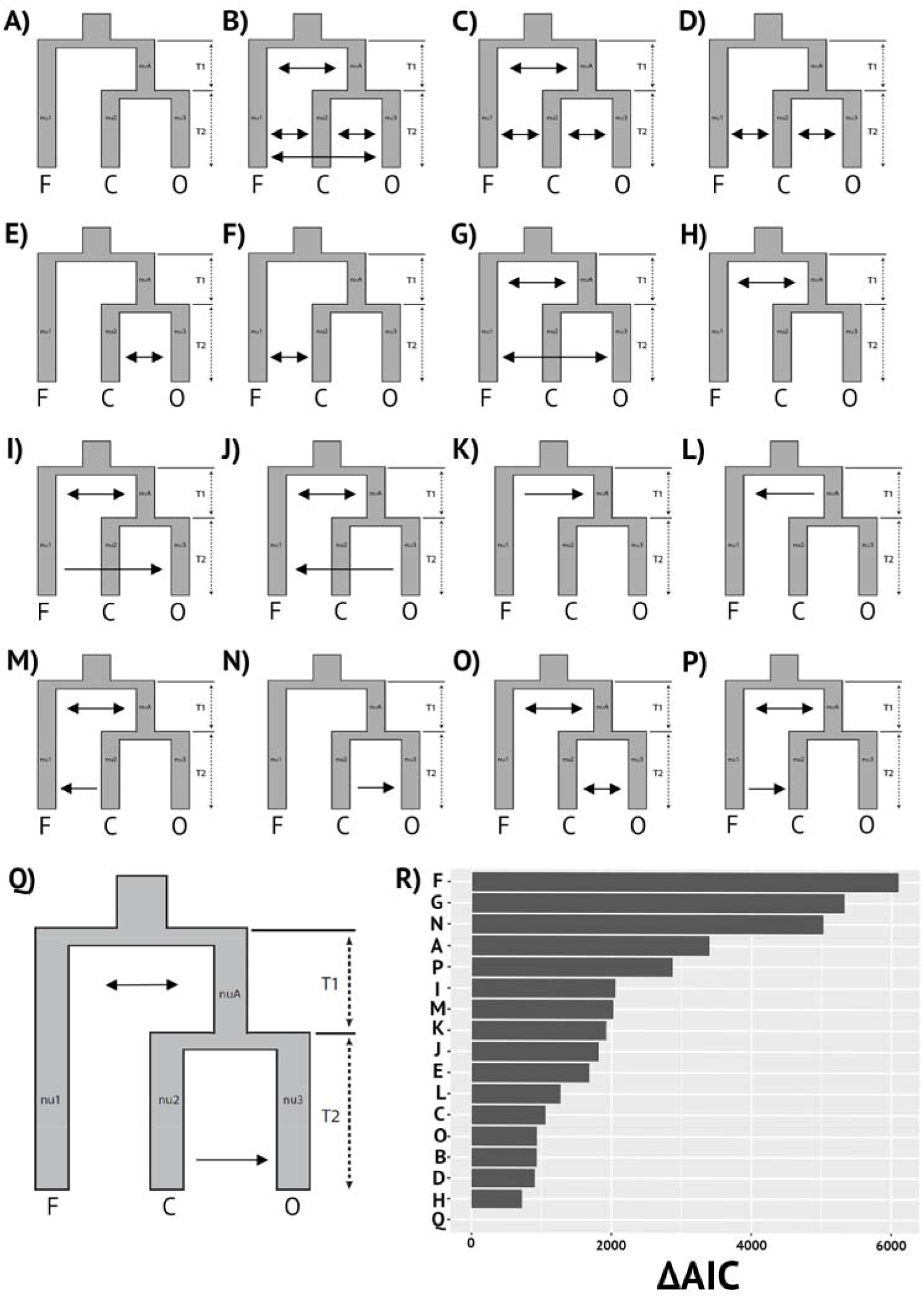
Schematic model summaries (A-Q) and results of demographic model selection analyses in *dadi* testing diversification scenarios in three cryptic species of *Entacmaea quadricolor* in the Japanese Archipelago. Each model is a three-population isolation-migration model with variation in the directionality (arrows) and timing (T1 & T2) of gene flow between co-occurring cryptic species. F = *Amphiprion frenatus* hosting *E. quadriolor* species in Mainland Japan, C = *A. clarkii* hosting *E. quadriolor* species in Mainland Japan, and O = *A. clarkii* hosting *E. quadricolor* species in the Ogasawara Islands (see Fig. 1). Model Q represents the best fit demographic model as determined by Akaike Information Criterion (AIC). R) ΔAIC results, which is the difference between the AIC scores of each model and the best fit model.

## Discussion

Our results unequivocally identify the bubble-tip sea anemone *Entacmaea quadricolor* as a diverse species complex shaped by both its mutualism with clownfishes and geographic distribution. Additionally, our data support the hypothesis that *E. quadricolor* may have undergone its own adaptive radiation. Our discovery that clownfishes can differentiate and ecologically segregate by co-occurring cryptic *E. quadricolor* lineages in Japan, and that there is an evolutionary basis for clownfish host specialist and generalist species of *E. quadricolor* on broader biogeographic scales, points to the importance of co-evolutionary processes generating and maintaining biodiversity in both the clownfishes and host sea anemones.

### Species delimitation and phylogeography of the Entacmaea quadricolor species complex

With the aid of independent genomic datasets we recover seven monophyletic lineages. We choose to conservatively delimit five cryptic species of *E. quadricolor*: I) Ogasawara Islands II) Ryukyu Islands and Mainland Japan (*A. clarkii*-hosting), III) Maldives, IV) Ryukyu Islands Japan + Singapore (*A. frenatus*-hosting), and V) Australia (Figure 3). For consistency and with future research in mind we elect to designate newly delimited species using Roman numerals (I-V; Figure 3). If additional evidence emergences that justifies further splitting our species here, a lowercase letter can be added to create a consistent alphanumeric system to informally recognize cryptic *E. quadricolor* species (e.g. Ia, IIa) until a formal revision can be completed. We currently elect not to further delimit *E. quadricolor sp.* IV (*A. frenatus*-hosting anemones from Japan and Singapore) or *E. quadricolor sp.* V (Great Barrier Reef and Solitary Islands – Australia) although we recovered these sample localities as monophyletic. Genetic divergences are not as deep as other splits in our phylogenetic reconstruction, and we do not have other additional lines of evidence to differentiate these groups (e.g. morphology/ecology), and so we elect not to delimit these as species for now (Figure 3).

The bubble-tip sea anemone has long been hypothesized to be a species complex, due largely to the highly variable phenotypic diversity encountered throughout its range (e.g. Titus et al. 2024). Traditional PCR-based sequencing hinted at the prospect of multiple species being present, even within Japan (Titus et al. 2019a) but the poorly resolving Sanger-loci were inconclusive. More recently, genomic approaches recovered similar genetic clusters within the Japanese Archipelago to ours here, yet lacked the biogeographic sampling needed to put this diversity into species-level context (Kashimoto et al. 2023). Our combined ddRADseq and UCE approach conclusively delimit three *E. quadricolor* species present within the Japanese Archipelago. We reveal that sympatric *E. quadricolor* diversity from the Ryukyu Islands are the result of geographic range overlap between two deeply diverged lineages (*E. quadricolor sp.* II and IV) that have evolved to host different clownfishes, rather than endemic *in situ* diversification within Japan. Subsequent allopatric speciation explains divergence between *E. quadricolor sp.* II (Ryukyu Islands and Mainland Japan) and *E. quadricolor sp.* I (Ogasawara Islands) that host *A. clarkii*. We make this interpretation as the *A. frenatus-*hosting *E. quadricolor sp.* IV from the Ryukyu Islands Japan is more closely related to individuals from Singapore than it is to the co-occurring *E. quadricolor sp.* II that hosts *A. clarkii*.

Demographic model selection in *dadi* demonstrates that ancestral divergence between the *A. frenatus* and *A. clarkii*-hosting species proceeded with bi-directional symmetric migration in T1. In T2, however, co-occurring *E. quadricolor sp.* II and IV lineages that host different species of clownfishes exist in complete genetic isolation with no signatures of contemporary migration—reinforcing species delimitation results that these are true biological species. Our best fit model also indicates that *E. quadricolor sp.* I from Ogasawara Islands diverged from *E. quadricolor sp.* II (Ryukyu Island and Mainland Japan) with unidirectional gene flow to Ogasawara. The Ogasawara Islands are located >1,000 km from Mainland Japan and separated by the Kuroshio Current, which is a deep fast-moving ocean current that flows north. Spin-off eddies, rotating clockwise and travelling east could potentially provide periodic larval transport to Ogasawara from Mainland Japan and explain the directionality of gene flow during diversification. The Japanese Archipelago has long been recognized as an important marine biogeographic region as it sits just north of the Coral Triangle where the Pacific and Indian Oceans meet (Motomura et al. 2007; Bowen et al. 2016; Endo et al. 2022). Our comprehensive fine-scale sampling and species delimitation analyses reinforces Japan as a marine biodiversity hotspot and disentangles the complex evolutionary and biogeographic processes that can contribute to generating biodiversity within this region (e.g. Bowen et al. 2016; Reimer et al. 2019).

Beyond Japan, hierarchical phylogenetic relationships among our newly delimited *E. quadricolor* species complex reveals the importance of broad-scale biogeographic processes that shapes diversity within this group. Strong geographic signal is evident within our bait-capture sequence data, and we recover most (but not all) sample localities as monophyletic. Even at the intraspecific levels we see signatures that demonstrate that the geographic scale of dispersal may be small for *E. quadricolor*. Previous work on *E. quadricolor* larval biology indicates planula larvae can metamorphose and settle as early as five days post fertilization, with most larvae settling within 10 days (Scott and Harrison 2008). Short larval durations could explain patterns in our data.

We recover intraspecific phylogeographic lineages within the *A. frenatus*-hosting *E. quadricolor sp.* IV, separating the Ryukyu Islands Japan from Singapore, and *E. quadricolor sp.* V, separating tropical (Great Barrier Reef) and subtropical (Solitary Islands) populations in Eastern Australia. Interestingly, geography has had a similar impact on the diversification of clownfishes (Litsios et al. 2012, 2014; Gaboriau et al. 2024). Many clownfish species have small range sizes that are highly partitioned geographically and with minimal overlap with sister taxa, particularly within the Coral Triangle (Litsios et al. 2012, 2014; Gaboriau et al. 2024). Dispersal kernels that have been estimated for some clownfish species have been shown to be quite small (<30km) within the Coral Triangle (e.g. Pinsky et al. 2010, 2017) and more extensive in regions outside the center of marine biodiversity (e.g. Simpson et al. 2014). The range sizes for our newly delimited *E. quadricolor* species are unknown until further fine-scale sampling and sequencing can be conducted in the Coral Triangle, but our findings indicate that geographic processes will continue to be key to fully understand diversification of this species complex.

### The Entacmaea quadricolor adaptive radiation?

Although clearly important, geography alone cannot fully explain diversification within the *E. quadricolor* species complex. Our data indicate that mutualism with clownfishes is also a key part of the evolutionary history of this species complex. Within the Japanese Archipelago we find that different species of clownfishes ecologically segregate between co-occurring cryptic hosts species (as did Kashimoto et al. 2023), but that this pattern is not simply the result of local processes unique to Japan. Remarkably, our bait-capture data show differentiation by cryptic host species is part of a deeper evolutionary signal within the *E. quadricolor* complex where some lineages serve as hosts to the host generalist clownfish species *A. clarkii*, which lives mutualistically on all 10 nominal clownfish-hosting anemone species, and others serve as host to the host specialist clownfish *A. frenatus*, which is only found with *E. quadricolor*.

Central to our interpretation of the evolutionary history of this species complex is the recognition that co-occurring *E. quadricolor* diversity within the Japanese Archipelago is not the result of endemic *in situ* diversification, as well as placement of the newly delimited species from the Maldives (*E. quadricolor sp.* III). Our phylogenetic analyses recover all *E. quadricolor* species that host *A. clarkii* belonging to a monophyletic clade. This clade includes *E. quadricolor sp.* I and II from Japan, which were recovered as allopatric sister species in all analyses, and *E. quadricolor sp.* III (Maldives), which also hosts *A. clarkii*. The hierarchical relationship linking Japan to the Maldives, to the exclusion of Singapore, runs counter to most phylogeographic patterns from the Indo-West Pacific. Typically, the deepest phylogeographic splits within tropical marine species complexes from this region partition Indian and Pacific Oceans into separate clades, with the Indo-Australian Archipelago (IAA) in the Coral Triangle serving as a well-resolved barrier (reviewed by Bowen et al. 2016). The presence of a deeper evolutionary signal within *E. quadricolor* connected to clownfish identity suggests the diversity of this species complex has an origin that is linked directly to the ecology of the mutualism. If so, our data may provide the first supporting evidence for the hypothesis that *E. quadricolor* has undergone its own adaptive radiation in response to mutualism with clownfishes.

Adaptive radiation has been defined as the “evolution of ecological and phenotypic diversity within a rapidly multiplying lineage” (Gavrilets and Vose 2005). Adaptive radiation requires a key innovation that provides a fitness advantage, the evolution of morphological and functional phenotypes that are linked to ecological niche space, and evidence of rapid speciation from a common ancestor (e.g Schluter 2020). To date, no key adaptive innovation has been proposed or identified for the clownfish-hosting sea anemones. We propose that mutualism with clownfishes should be considered a key adaptive innovation for host sea anemones. The mutualistic benefits provided to sea anemones by hosting clownfishes are well established, and broadly, the presence of clownfishes is necessary for host anemone survival on tropical coral reefs. Clownfishes provide novel sources of nitrogen via waste byproducts that are taken up by the host anemones and their endosymbiotic algae— providing increased protein synthesis and algal density and allowing host anemones to thrive on oligotrophic coral reefs (Roopin et al. 2008; Roopin and Chadwick 2009). Clownfishes also facilitate gas and nutrient transfer as they move through their host anemone tentacles (Szczebak et al. 2013; Herbert et al. 2017) and provide reciprocal protection by defending their sea anemone hosts from butterflyfishes, turtles, and other predators (Fautin 1991; Godwin and Fautin 1992). Sea anemones with clownfish symbionts have increased growth and survivorship over those that do not (Porat and Chadwick-Furman 2004, 2005). Finally, the clownfish-hosting anemones have historically been labeled “giant” tropical sea anemones because they attain large size classes—some reaching 1m in diameter (e.g. Fautin and Allen 1992). No other tropical sea anemones reach the same sizes and occur as free-living individuals on Indo-Pacific coral reefs without clownfish symbionts. Taken together, the host anemones are likely able to compete for, and maintain, habitat space on densely populated coral reef ecosystems because of their mutualism with clownfishes.

If mutualism with clownfishes is adaptive, our findings for *E. quadricolor* begin to fulfill other requirements of adaptive radiation. The discovery of an evolutionary signal linking generalist and specialist clownfish species to cryptic *E. quadricolor* speciation indicates functional and ecological differences between co-occurring host species that would limit niche overlap and competition for resources. Our phenotypic/morphological data reveals previously unrecognized phenotypic differences between cryptic hosts species that are linked to clownfish symbiont identity. Historically, sea anemone color and pattern has been held as an uninformative taxonomic character. None of the phenotypic characters we explored here could be considered synapomorphic, yet we do find that tentacle tip shape and tentacle tip pattern are strongly predictive of cryptic host species in Japan. Tentacles with blunt-rounded bubble tips were exclusively found in *E. quadricolor sp.* IV which hosted *A. frenatus,* while tentacles with elongated tips extending from the bulbous swelling were nearly exclusively found in both *E. quadricolor sp.* I and II which hosted *A. clarkii*. Tentacle tip pattern also was strongly predictive of cryptic species, with *E. quadricolor sp.* IV anemones having a dull/matte tentacle tip pattern and with *E. quadricolor sp.* I and II anemones having striated patterns and pink tips. Interestingly, these phenotypes may have an evolutionary signal consistent across *E. quadricolor* lineages. While we did not have the images to conduct similar phenotypic analyses elsewhere, representative images from Singapore (*A. frenatus*-hosting *E. quadricolor sp.* IV) and the Maldives (*A. clarkii*-hosting *E. quadricolor sp.* III) align well with the phenotypes we recovered from Japan for both specialist and generalist *E. quadricolor* species (Figure 3d & 3e).

Finally, our phylogenetic reconstruction and divergence time analyses demonstrate that the *E. quadricolor* species complex has diversified rapidly from a common ancestor to at least five unique species within the last 4-5 million years. Our species delimitation erred on being conservative to not over-delimit species, but our phylogenetic analyses delimited seven monophyletic lineages. Our study also focused on fine-scale sampling within the Japanese Archipelago and adjacent biogeographic regions. We have yet to conduct comprehensive sampling and sequencing to include the entire biogeographic range of the *E. quadricolor* complex, which extends from the Northern Red Sea and Arabian Peninsula, the Western Indian Ocean, Western Australia, Coral Triangle, and well into the Central Pacific Ocean. Additional cryptic species undoubtedly remain to be delimited as the true scope of diversification within the *E. quadricolor* complex comes into focus. In sum, our data fail to reject the hypothesis that *E. quadricolor* has undergone an adaptive radiation in response to its mutualism with clownfishes. Whether mutualism with clownfishes is a true key innovation, and the *E. quadricolor* complex exhibits enough ecological niche partitioning and character displacement to be considered an adaptive radiation, is unclear but will remain an important future hypothesis to test.

### Emerging patterns of co-diversification within the clownfish-sea anemone symbiosis

Mutualism establishes a strong *a priori* hypothesis for co-evolutionary patterns between partner symbionts. Yet within the clownfish-sea anemone symbiosis co-evolutionary patterns have remained elusive. Unlike the clownfishes, which have descended from a common ancestor and belong to the same genus, the 10 nominal species of host sea anemones belong to five genera within three clades that have evolved symbiosis with clownfishes independently. Thus, no taxonomic justification for classic co-diversification/cladogenesis existed based on traditional sea anemone systematics (Fautin and Allen 1992; Titus et al. 2019, 2024; De Jode et al. 2024), and previous efforts with traditional PCR-based genetic markers have been unsuccessful (Ngyuyen et al. 2020). Recently, however, the first divergence time estimates for the clownfish-hosting sea anemones using genomic approaches revealed broadly coincident diversification times between sea anemones and the clownfish-radiation (De Jode et al. 2024), providing the first evidence that both symbiotic partners were diversifying around the same time. Here, our data reveal an evolutionary basis for specialist and generalist lineages of host sea anemones within just one host species complex. Our data indicate that clownfishes are not merely settling in locally available hosts but recruiting to specialized host lineages with which they have co-evolved. The discovery of what appears to be clownfish host generalist and clownfish host specialist *E. quadricolor* lineages represents a major insight that furthers our evolutionary understanding of the clownfish-sea anemone symbiosis and underscores the importance of disentangling the systematics and diversity of the host sea anemones for a comprehensive understanding of this charismatic mutualism.

## Acknowledgements

We thank Laura Simmons and Cairns Marine Inc. for providing tissue samples of *Entacmaea quadricolor* from the Great Barrier Reef (Australia). Dr. Christopher Meyer (National Museum of Natural History) and Small Island Lodge are thanked for fieldwork and logistics in the Maldives. Charlotte Benedict and Robert Laroche assisted with DNA extractions for all samples at the American Museum of Natural History and were funded by a National Science Foundation (NSF) Research Experience for Undergraduate award DBI-1358465 to M. Siddall. Field research and genomic sequencing were funded by NSF awards DEB-1934274 to BMT and ER, and DEB-145781 to ER. Field research in Japan was funded by the Japan Society for the Promotion of Science (JSPS) Kakenhi Grants (JP255440221 to KY, and JP17K15198, JP17H01913, and 23K21774 grants to TF), and Kagoshima University (Establishment of Research and Education Network of Biodiversity and its Conservation in the Satsunan Islands project to TF). Further funding to BMT was provided by University of Alabama start-up research funds, and through a Gerstner Scholar Postdoctoral Fellowship and Gerstner Family Foundation, The Lerner Gray Fund for Marine Research, and Richard Gilder Graduate School at the American Museum of Natural History.

