## Supplemental Material for "Nemo knows: clownfishes differentiate cryptic host species across fine and broad geographic scales and reveal a potential adaptive radiation in the clownfish-hosting sea anemones"

**Supplementary Methods**

### *Sample collection*

To search for patterns of cryptic species-level diversity within *Entacmaea quadricolor*, we first conducted fine-scale phylogeographic surveys and sampling across the Japanese Archipelago. We surveyed and photographed N = 126 individuals, and collected N = 93 tentacle clippings from 38 sample localities, including the remote Ogasawara Islands ~1000 km from Mainland Japan (Figure 1; Table S1). Samples were collected by hand using SCUBA between 1-29 m depth. *In situ* photos were taken of each anemone, clownfish symbionts were identified and quantified, and one or two tentacles were sampled using forceps and placed into individually labeled collection bags. On shore, samples were preserved in 95% ethanol. To place our Japanese samples into a broader biogeographic context, we also collected *E. quadricolor* samples from adjacent biogeographic regions including Singapore, Maldives, Australia, and the Philippines (Table S3). Samples were collected and preserved as above.

### *DNA Extraction, Library Preparation, and Sequencing*

Following collection, total genomic DNA was extracted using a modified DNeasy Blood and Tissue Kits (QIAGEN Inc.) at the American Museum of Natural History and Dauphin Island Sea Lab. DNA was eluted from spin columns twice using 80µl of Buffer AE warmed to 50℃. From the resulting DNA extractions, 5µl were loaded on a 1% agarose gel alongside a 5 kb ladder to confirm the DNA was high molecular weight (>1000kb fragment length). Samples were quantified using a Qubit 4 Fluorometer (Life Technologies) and standardized to a concentration of 25ng/µl for library preparation and sequencing. Samples below the concentration threshold were vacuum centrifuged until they met the cutoff.

To test for undescribed cryptic diversity within *E. quadricolor* and place the resulting diversity into broader phylogeographic context, we generated two independent genomic datasets. First, for all samples we collected throughout the Japanese Archipelago, we used a double-digest restriction-site associated DNA sequencing (ddRADseq) approach for cryptic species discovery [56]. This method generates thousands of unlinked single-nucleotide polymorphisms (SNPs) across the genome per individual and is particularly cost effective when no genomic resources exist for the focal taxa and samples sizes are large. Within anthozoans, RADseq has been used effectively to delimit species in corals [e.g. 57-59], and within Actiniaria, several studies have been used for assessing phylogeographic variation and cryptic species delimitation [60-65]. Further, within tropical anthozoans that harbor endosymbionts, given a sufficient sample size and broad sampling distributions, ddRADseq is a robust and appropriate sequencing approach as endosymbiont sequence reads are largely filtered out during the dataset assembly pipeline [63].

Library preparation and ddRAD sequencing were conducted at the University of Wisconsin Biotechnology Center DNA Sequencing Facility. Prior to library preparation, multiple optimization steps were completed to determine NSIL and Bfal as the best restriction enzyme combination for *E. quadricolor.* Following digestion, fragment sizes between 400-800 bp were selected for library preparation. Samples were barcoded with Illumina compatible barcode adapters, pooled, and sequenced on an Illumina NovaSeqX+ using 150bp paired-end reads.

Next, we used bait-capture sequencing targeting ultra-conserved elements (UCEs) and exon loci to produce a second genomic dataset and place the resulting cryptic lineages recovered from Japan into hierarchical biogeographic context. Accordingly, we tested whether the *E. quadricolor* lineages in Japan were diversifying *in situ* or were the result of geographical range overlap of more distantly related species. We re-sequenced N = 64 individuals from Japan along with N = 46 additional samples from adjacent biogeographic regions including Singapore [66], Philippines, Australia, and Maldives (Table S2). We used this alternative genomic approach as *E. quadricolor* does not have a closely related sister species that would be suitable to use as an outgroup taxon to root a ddRADseq dataset [23, 24] and previously published bait-capture data for sea anemones are publicly available on GenBank (Table S3) [34, 35].

Bait-capture libraries were prepared at both Harvey Mudd College and Arbor BioSciences (Ann Arbor, MI) following the protocol developed by Quattrini et al. [34]. We used the Hexacorallia specific bait set from Cowman et al. [67], which was further modified to specifically target actiniarians [61]. The resulting bait set contained 17,268 baits targeting 2496 loci (e.g. exons and UCEs), which were synthesized by Arbor BioSciences. For each sample, up to 1,000ng of genomic DNA was carried forward for library preparation, which was sheared to a target fragment size of 400-800 bp using a Covaris Ultrasonicator. Library preparation was performed using a Kapa Hyper Prep Kit for bait-capture sequencing, with universal Y-yoke oligonucleotide adapters and iTru dual-indexed primers [68]. Libraries were pooled into equimolar ratios (100 ng) and target enrichment was performed using MyBaits v.IV protocol with a 500 ng/rxn concentration of baits [61]. Bait-capture enriched libraries were sequenced at Arbor BioSciences on an Illumina NovaSeq using 150 bp paired-end sequencing. Samples were sequenced across three individual Illumina runs.

### *Dataset assembly*

Following sequencing, raw ddRADseq data were demultiplexed, aligned, and assembled *de novo* using ipyrad [69]. We set the clustering threshold (Wclust) to 0.90 to assemble reads into loci and sequencing depth (e.g. mindepth_statistical and mindepth_majrule) parameters were set to 10 to reduce heterozygous calls due to sequencing error. The min_samples_locus was set to 76 as that would represent approximately 75% occupancy across samples for the locus to be retained in the ddRADseq dataset. One SNP per locus was randomly selected to build an unlinked SNP dataset. Raw ddRADseq data were deposited at the NCBI Sequence Read Archive under BioProject XXXXX (Table S1).

To assemble UCE and exon loci from raw bait-capture sequence data we used the program PHYLUCE [68] and the hexa_v2_final bait set [61, 67]. After demultiplexing, raw sequences for each individual sample were cleaned and trimmed using illumiprocessor [70, 71]. Cleaned sequences were then assembled into contigs using SPAdes v3.14.1 [72] with the -careful and -cov-cutoff 2 parameters. We then used PHYLUCE, as described in online tutorials to extract exon and UCE loci and assemble the final dataset. UCE baits were matched and then extracted using phyluce_assembly_ match_contigs_probes and phyluce_assembly_get_match_counts. We created a final dataset with 75% completeness (i.e. no more than 25% missing data at a single locus). Our dataset was aligned using MAFFT [73] and internally trimmed using the version of Gblocks inside PHYLUCE (phyluce_align_get_gblocks_trimmed_alignments_from_untrimmed) with the default parameters. For downstream bait-capture analyses, *Epiactis georgiana* Carlgren, 1927 was used as an outgroup species as this is currently the most closely related known species to *E. quadricolor* [24].

### *Fine scale phylogeography and species delimitation in the Japanese Archipelago*

To test whether there is evidence that *E. quadricolor* is a cryptic species complex within the Japanese Archipelago we used our fine-scale ddRADseq dataset to conduct three species discovery analyses. We first conducted a principal coordinates analysis (PCA) as a preliminary investigation to visualize the genetic variation present in the Japanese Archipelago in our unlinked SNP dataset generated by ipyrad. Preliminary PCAs revealed three genetic clusters, two of which were co-occurring and linked to clownfish identity. However, the *A. frenatus*-hosting cluster had low sample sizes. Some originally sequenced *A. frenatus*-hosting samples did not make it through preliminary dataset assembly steps since they failed to meet the 75% missing data threshold. Thus, in order to minimize the amount of missing data (NAs) for our PCA while also retaining the most individuals from our groups of interest, the sample with the greatest number of missing loci that belonged to the *A. frenatus*-hosting anemone lineage was used to select which loci would be used for the PCA matrix. Of the resulting matrix, only samples which had at minimum of 60% SNP occupancy were retained. Remaining NAs were replaced with the mean value and the final PCA was calculated using the ade4 package in R v4.3.1 [74, 75].

We next performed a Discriminant Analysis of Principal Components (DAPC) on the ddRADseq data to assign individuals of *E. quadricolor* from Japan to genetic clusters using the R package adegenet [76]. The optimal value of *k* was obtained using a *k*-means clustering algorithm (find.clusters()) and the *k* value with the lowest BIC value was selected. To determine the number of principal components to retain for the DAPC, an ⍺-score optimization run was done and cross validated by selecting the number of PCs with the highest average prediction success per group. A separate DAPC analysis was conducted for *k* = 2-5 to compare group assignments of different *k* values to our biological lineages. Finally, we used the Bayesian genetic clustering approach implemented in the program fastStructure [77] to delimit major genetic clusters and assign individuals to putative species. Using default parameters, we tested *k* values from 2 to 5 to determine number of genetic clusters that maximized the marginal likelihood for our dataset. All clustering programs identified the same three major genetic clusters, and we computed pairwise F_ST_ to determine the degree of genetic differentiation within and between clusters using the R package hierfstat [78].

### *Demographic modeling*

We conducted demographic modeling in *dadi* (diffusion approximations for demographic inference [91]) to test competing diversification scenarios involving our newly delimited *E. quadricolor* from the Japanese Archipelago (*A. frenatus*-hosting, *A. clarkii*-hosting, and Ogasawara Islands). We tested whether ancestral diversification between *A. frenatus* and *A. clarkii* lineages occurred with or without geneflow, as well as tested whether co-occurring lineages in Japan were reproductively isolated. We built a set of 17 demographic models (Figure 4) following protocols and pipelines detailed by [92]. Each model was a three-population isolation-migration model that varied in the directionality and timing of gene flow between *E. quadricolor* species.

To model the evolutionary history of *E. quadricolor*, we built a three-dimensional Joint-folded Site Frequency Spectrum (JSFS) using our ddRADseq dataset from the Japanese Archipelago. Individual anemones were assigned to putative species based on species delimitation results. All loci not in Hardy-Weinberg Equilibrium (p > 0.05) were removed using VCFTools [93]. Model simulations were conducted with consecutive rounds of optimization, where multiple replicates and previous parameter estimates from best scoring replicates were used to seed subsequent simulations. The default dadi_pipeline settings were used for each round (replicates = 10, 20, 30, 40; maxiter = 3, 5, 10, 15; fold = 3, 2, 2, 1) and the parameter optimization followed the Nelder-Mead method (optimize_log_fmin). The optimized parameters of each replicate were used to simulate the 3D-JSFS and estimate the log-likelihood of the JSFS of the model. The best fit demographic model was selected using Akaike Information Criterion and model probabilities were calculated following Burnham and Anderson [94].

**Supplementary Tables**

**Table S1.** List of individual bubble-tip sea anemones, *Entacmaea* quadricolor, collected across the Japanese Archipelago and sequenced using double digest Restriction Site Associated DNA sequencing. Information includes Sample ID, sample Prefecture, Latitude and Longitude of sample locality, Depth (meters), clownfish species identification, and GenBank Accession Number.

| **Sample ID** | **Anemone species** | **Prefecture** | **Latitude (N)** | **Longitude (E)** | **Depth (m)** | **Clownfish sp.** | **Accession #** |
| --- | --- | --- | --- | --- | --- | --- | --- |
| K008 | *Entacmaea quadricolor* | Okinawa | 26.441666 | 127.712222 | 4.5 | *Amphiprion clarkii* | TBD |
| K016 | *Entacmaea quadricolor* | Okinawa | 26.537777 | 128.079722 | 6 | *Amphiprion frenatus* | TBD |
| K017 | *Entacmaea quadricolor* | Okinawa | 26.537777 | 128.079722 | 5.5 | *Amphiprion frenatus* | TBD |
| K019 | *Entacmaea quadricolor* | Shizuoka | 35.02164 | 138.86691 | 1 | None | TBD |
| K039 | *Entacmaea quadricolor* | Kagoshima | 28.14976 | 129.25402 | 4.3 | *Amphiprion frenatus* | TBD |
| K040 | *Entacmaea quadricolor* | Kagoshima | 28.14976 | 129.25402 | 6.2 | *Amphiprion frenatus* | TBD |
| K041 | *Entacmaea quadricolor* | Kagoshima | 28.40708 | 129.45211 | 8.7 | *Amphiprion frenatus* | TBD |
| K042 | *Entacmaea quadricolor* | Kagoshima | 28.14976 | 129.25402 | 4.7 | *Amphiprion clarkii* | TBD |
| K044 | *Entacmaea quadricolor* | Kagoshima | 28.1736 | 129.27993 | 6.7 | *Amphiprion frenatus* | TBD |
| K055 | *Entacmaea quadricolor* | Kagoshima | 28.292837 | 129.475717 | 6.5 | *Amphiprion frenatus* | TBD |
| K062 | *Entacmaea quadricolor* | Kagoshima | 29.858647 | 129.833992 | 10 | None | TBD |
| K063 | *Entacmaea quadricolor* | Kagoshima | 31.252073 | 130.374979 | N/A | None | TBD |
| K064 | *Entacmaea quadricolor* | Kagoshima | 31.549561 | 130.647044 | 6 | *Amphiprion clarkii* | TBD |
| K071 | *Entacmaea quadricolor* | Kagoshima | 31.549561 | 130.647044 | 10 | *Amphiprion clarkii* | TBD |
| K072 | *Entacmaea quadricolor* | Kanagawa | 35.294189 | 139.556109 | 2 | None | TBD |
| K074 | *Entacmaea quadricolor* | Kanagawa | 35.294189 | 139.556109 | 2 | None | TBD |
| K078 | *Entacmaea quadricolor* | Wakayama | 35.454167 | 135.748611 | 20 | *Amphiprion clarkii* | TBD |
| K079 | *Entacmaea quadricolor* | Wakayama | 35.454167 | 135.748611 | 20 | *Amphiprion clarkii* | TBD |
| K082 | *Entacmaea quadricolor* | Wakayama | 33.476389 | 135.747222 | 10 | *Amphiprion clarkii* | TBD |
| K084 | *Entacmaea quadricolor* | Wakayama | 33.476389 | 135.747222 | 10 | *Amphiprion clarkii* | TBD |
| K085 | *Entacmaea quadricolor* | Wakayama | 33.476389 | 135.747222 | 12 | *Amphiprion clarkii* | TBD |
| K089 | *Entacmaea quadricolor* | Kagoshima | 30.462636 | 130.500703 | 15 | *Amphiprion clarkii* | TBD |
| K094 | *Entacmaea quadricolor* | Kagoshima | 30.464414 | 130.499839 | 18 | *Amphiprion clarkii* | TBD |
| K097 | *Entacmaea quadricolor* | Kagoshima | 30.464414 | 130.499839 | 12 | *Amphiprion clarkii* | TBD |
| K098 | *Entacmaea quadricolor* | Kagoshima | 30.462636 | 130.500703 | 19.7 | *Amphiprion clarkii* | TBD |
| K107 | *Entacmaea quadricolor* | Shizuoka | 34.84619 | 138.765084 | intertidal | None | TBD |
| K108 | *Entacmaea quadricolor* | Shizuoka | 34.84619 | 138.765084 | intertidal | None | TBD |
| K109 | *Entacmaea quadricolor* | Shizuoka | 34.84619 | 138.765084 | intertidal | None | TBD |
| K117 | *Entacmaea quadricolor* | Kagoshima | 30.814222 | 131.029028 | 21 | *Amphiprion clarkii* | TBD |
| K118 | *Entacmaea quadricolor* | Kagoshima | 30.814222 | 131.029028 | 21 | *Amphiprion clarkii* | TBD |
| K119 | *Entacmaea quadricolor* | Kagoshima | 30.814222 | 131.029028 | 21 | *Amphiprion clarkii* | TBD |
| K152 | *Entacmaea quadricolor* | Ogasawara | 27.1208333 | 142.199222 | 15.4 | *Amphiprion clarkii* | TBD |
| K153 | *Entacmaea quadricolor* | Ogasawara | 27.1208333 | 142.199222 | 16.9 | *Amphiprion clarkii* | TBD |
| K155 | *Entacmaea quadricolor* | Ogasawara | 27.1012222 | 142.235528 | 18.1 | *Amphiprion clarkii* | TBD |
| K156 | *Entacmaea quadricolor* | Ogasawara | 27.1012222 | 142.235528 | 17.8 | *Amphiprion clarkii* | TBD |
| K157 | *Entacmaea quadricolor* | Ogasawara | 27.1211111 | 142.1805 | 27 | *Amphiprion clarkii* | TBD |
| K158 | *Entacmaea quadricolor* | Ogasawara | 27.1211111 | 142.1805 | 29 | *Amphiprion clarkii* | TBD |
| K159 | *Entacmaea quadricolor* | Ogasawara | 27.1211111 | 142.1805 | 8 | *Amphiprion clarkii* | TBD |
| K160 | *Entacmaea quadricolor* | Ogasawara | 27.1211111 | 142.1805 | 24.7 | *Amphiprion clarkii* | TBD |
| K161 | *Entacmaea quadricolor* | Ogasawara | 27.1211111 | 142.1805 | 21.4 | *Amphiprion clarkii* | TBD |
| K162 | *Entacmaea quadricolor* | Ogasawara | 27.1211111 | 142.1805 | 19.3 | *Amphiprion clarkii* | TBD |
| K163 | *Entacmaea quadricolor* | Ogasawara | 27.1211111 | 142.1805 | 22 | *Amphiprion clarkii* | TBD |
| K164 | *Entacmaea quadricolor* | Ogasawara | 27.1211111 | 142.1805 | 10 | *Amphiprion clarkii* | TBD |
| K165 | *Entacmaea quadricolor* | Ogasawara | 27.0814167 | 142.187389 | 17 | *Amphiprion clarkii* | TBD |
| K167 | *Entacmaea quadricolor* | Ogasawara | 27.0814167 | 142.187389 | 11.1 | *Amphiprion clarkii* | TBD |
| K168 | *Entacmaea quadricolor* | Ogasawara | 27.0814167 | 142.187389 | 18.1 | *Amphiprion clarkii* | TBD |
| K169 | *Entacmaea quadricolor* | Ogasawara | 27.12025 | 142.162972 | 15.7 | *Amphiprion clarkii* | TBD |
| K170 | *Entacmaea quadricolor* | Ogasawara | 27.12025 | 142.162972 | 9.6 | *Amphiprion clarkii* | TBD |
| K171 | *Entacmaea quadricolor* | Ogasawara | 27.12025 | 142.162972 | 23.2 | *Amphiprion clarkii* | TBD |
| K173 | *Entacmaea quadricolor* | Ogasawara | 27.12025 | 142.162972 | 14.8 | *Amphiprion clarkii* | TBD |
| K176 | *Entacmaea quadricolor* | Ogasawara | 27.12025 | 142.162972 | 20 | *Amphiprion clarkii* | TBD |
| K177 | *Entacmaea quadricolor* | Ogasawara | 27.0413333 | 142.202611 | 16 | *Amphiprion clarkii* | TBD |
| K178 | *Entacmaea quadricolor* | Ogasawara | 27.0413333 | 142.202611 | 22 | *Amphiprion clarkii* | TBD |
| K179 | *Entacmaea quadricolor* | Ogasawara | 27.0413333 | 142.202611 | 6.8 | *Amphiprion clarkii* | TBD |
| K180 | *Entacmaea quadricolor* | Ogasawara | 27.0413333 | 142.202611 | 8 | *Amphiprion clarkii* | TBD |
| K181 | *Entacmaea quadricolor* | Ogasawara | 27.0413333 | 142.202611 | 13 | *Amphiprion clarkii* | TBD |
| K182 | *Entacmaea quadricolor* | Ogasawara | 27.0413333 | 142.202611 | 7.4 | *Amphiprion clarkii* | TBD |
| K183 | *Entacmaea quadricolor* | Ogasawara | 27.0413333 | 142.202611 | 12.9 | *Amphiprion clarkii* | TBD |
| K184 | *Entacmaea quadricolor* | Ogasawara | 27.0413333 | 142.202611 | 13.5 | *Amphiprion clarkii* | TBD |
| K185 | *Entacmaea quadricolor* | Ogasawara | 27.0413333 | 142.202611 | 9.8 | *Amphiprion clarkii* | TBD |
| K186 | *Entacmaea quadricolor* | Ogasawara | 27.05425 | 142.176306 | 18 | *Amphiprion clarkii* | TBD |
| K187 | *Entacmaea quadricolor* | Ogasawara | 27.05425 | 142.176306 | 10 | *Amphiprion clarkii* | TBD |
| K188 | *Entacmaea quadricolor* | Ogasawara | 27.05425 | 142.176306 | 17.6 | *Amphiprion clarkii* | TBD |
| K189 | *Entacmaea quadricolor* | Ogasawara | 27.1186667 | 142.2295 | 27.2 | *Amphiprion clarkii* | TBD |
| K190 | *Entacmaea quadricolor* | Ogasawara | 27.1186667 | 142.2295 | 19.1 | *Amphiprion clarkii* | TBD |
| K191 | *Entacmaea quadricolor* | Ogasawara | 27.1186667 | 142.2295 | 15.5 | *Amphiprion clarkii* | TBD |
| K192 | *Entacmaea quadricolor* | Ogasawara | 27.1186667 | 142.2295 | 25.3 | *Amphiprion clarkii* | TBD |
| K193 | *Entacmaea quadricolor* | Ogasawara | 27.1186667 | 142.2295 | 23.8 | *Amphiprion clarkii* | TBD |
| K194 | *Entacmaea quadricolor* | Ogasawara | 27.1309444 | 142.221278 | 22 | *Amphiprion clarkii* | TBD |
| K195 | *Entacmaea quadricolor* | Ogasawara | 27.1309444 | 142.221278 | 25.2 | *Amphiprion clarkii* | TBD |
| K196 | *Entacmaea quadricolor* | Ogasawara | 27.1309444 | 142.221278 | 15.7 | *Amphiprion clarkii* | TBD |
| K197 | *Entacmaea quadricolor* | Ogasawara | 27.1309444 | 142.221278 | 23.2 | *Amphiprion clarkii* | TBD |
| K198 | *Entacmaea quadricolor* | Ogasawara | 27.1180556 | 142.173556 | 11.4 | *Amphiprion clarkii* | TBD |
| K199 | *Entacmaea quadricolor* | Ogasawara | 27.1833611 | 142.184917 | 18.5 | *Amphiprion clarkii* | TBD |
| K200 | *Entacmaea quadricolor* | Ogasawara | 27.1833611 | 142.184917 | 14.1 | *Amphiprion clarkii* | TBD |
| K201 | *Entacmaea quadricolor* | Ogasawara | 27.1833611 | 142.184917 | 14.3 | *Amphiprion clarkii* | TBD |
| K202 | *Entacmaea quadricolor* | Ogasawara | 27.1833611 | 142.184917 | 12.9 | *Amphiprion clarkii* | TBD |
| K203 | *Entacmaea quadricolor* | Ogasawara | 27.1833611 | 142.184917 | 17.5 | *Amphiprion clarkii* | TBD |
| K204 | *Entacmaea quadricolor* | Ogasawara | 27.1833611 | 142.184917 | 9.5 | *Amphiprion clarkii* | TBD |
| K205 | *Entacmaea quadricolor* | Ogasawara | 27.1833611 | 142.184917 | 16.7 | *Amphiprion clarkii* | TBD |
| K206 | *Entacmaea quadricolor* | Ogasawara | 27.1833611 | 142.184917 | 15.6 | *Amphiprion clarkii* | TBD |
| K207 | *Entacmaea quadricolor* | Kagoshima | 31.6244111 | 130.683233 | 11 | *Amphiprion clarkii* | TBD |
| K217 | *Entacmaea quadricolor* | Kochi | 32.774655 | 132.625274 | 22.3 | *Amphiprion clarkii* | TBD |
| K221 | *Entacmaea quadricolor* | Kochi | 32.768892 | 132.639893 | 1 | None | TBD |
| K222 | *Entacmaea quadricolor* | Kochi | 32.768892 | 132.639893 | 1 | None | TBD |
| K227 | *Entacmaea quadricolor* | Kochi | 32.856044 | 132.662497 | 3 | *Amphiprion clarkii* | TBD |
| K234 | *Entacmaea quadricolor* | Kochi | 32.753242 | 132.549453 | 15 | *Amphiprion clarkii* | TBD |
| K236 | *Entacmaea quadricolor* | Kochi | 32.761058 | 132.635619 | 25.1 | *Amphiprion clarkii* | TBD |
| K237 | *Entacmaea quadricolor* | Kochi | 32.761058 | 132.635619 | 21.3 | *Amphiprion clarkii* | TBD |
| K238 | *Entacmaea quadricolor* | Kochi | 32.774655 | 132.625274 | 13.2 | *Amphiprion clarkii* | TBD |
| K241 | *Entacmaea quadricolor* | Oita | 32.727222 | 131.921944 | 9.5 | *Amphiprion clarkii* | TBD |
| K242 | *Entacmaea quadricolor* | Oita | 32.727222 | 131.921944 | 10.6 | *Amphiprion clarkii* | TBD |
| K243 | *Entacmaea quadricolor* | Oita | 32.727222 | 131.921944 | 9.1 | *Amphiprion clarkii* | TBD |

**Table S2.** List of individual bubble-tip sea anemones, *Entacmaea quadricolor,* collected and sequenced using bait-capture sequencing targeting Ultra Conserved Elements (UCEs) and exon loci. Samples include individuals collected from Australia, Japan, Maldives, and Singapore. Information includes Sample ID, Country, Locality, Latitude and Longitude of sample locality, Depth (meters), clownfish species identification, and GenBank Accession Number.

| **Sample ID** | **Country** | **Locality** | **Latitude** | **Longitude** | **Depth (m)** | **Clownfish sp.** | **Accession #** |
| --- | --- | --- | --- | --- | --- | --- | --- |
| AUCSEQ7 | Australia | Coral Sea-Holmes Reef | 16.4500°S | 148.0000°E | N/A | N/A | TBD |
| AUGBREQ1 | Australia | Northern Great Barrier Reef | 15.872°S | 145.79935°E | N/A | N/A | TBD |
| AUGBREQ2 | Australia | Northern Great Barrier Reef | 15.872°S | 145.79935°E | N/A | N/A | TBD |
| AUGBREQ3 | Australia | Northern Great Barrier Reef | 15.872°S | 145.79935°E | N/A | N/A | TBD |
| AUGBREQ4 | Australia | Northern Great Barrier Reef | 15.872°S | 145.79935°E | N/A | N/A | TBD |
| AUGBREQ6 | Australia | Northern Great Barrier Reef | 15.872°S | 145.79935°E | N/A | N/A | TBD |
| EQ11GBR | Australia | Northern Great Barrier Reef | 15.872°S | 145.79935°E | N/A | N/A | TBD |
| EQ12GBR | Australia | Northern Great Barrier Reef | 15.872°S | 145.79935°E | N/A | N/A | TBD |
| EQ13GBR | Australia | Northern Great Barrier Reef | 15.872°S | 145.79935°E | N/A | N/A | TBD |
| EQ10AUSI | Australia | Solitary Islands | 29.92456°S | 153.3906°E | N/A | *Amphiprion akindynos* | TBD |
| EQ1SI | Australia | Solitary Islands | 29.92456°S | 153.3906°E | N/A | *Amphiprion akindynos* | TBD |
| EQ2SI | Australia | Solitary Islands | 29.92456°S | 153.3906°E | N/A | *Amphiprion akindynos* | TBD |
| EQ4SI | Australia | Solitary Islands | 29.92456°S | 153.3906°E | N/A | *Amphiprion akindynos* | TBD |
| EQ5AUSI | Australia | Solitary Islands | 29.92456°S | 153.3906°E | N/A | *Amphiprion akindynos* | TBD |
| EQ6AUSI | Australia | Solitary Islands | 29.92456°S | 153.3906°E | N/A | *Amphiprion akindynos* | TBD |
| EQ7AUSI | Australia | Solitary Islands | 29.92456°S | 153.3906°E | N/A | *Amphiprion akindynos* | TBD |
| EQ9AUSI | Australia | Solitary Islands | 29.92456°S | 153.3906°E | N/A | *Amphiprion akindynos* | TBD |
| K040 | Japan | Kagoshima | 28.1497°N | 129.25402°E | 6.2 | *Amphiprion frenatus* | TBD |
| K042 | Japan | Kagoshima | 28.1497°N | 129.25402°E | 4.7 | *Amphiprion clarkii* | TBD |
| K044 | Japan | Kagoshima | 28.1736°N | 129.27993°E | 6.7 | *Amphiprion frenatus* | TBD |
| K045EQ | Japan | Kagoshima | 28.4049°N | 129.452296°E | 15 | *Amphiprion frenatus* | TBD |
| K055 | Japan | Kagoshima | 28.2928°N | 129.475717°E | 6.5 | *Amphiprion frenatus* | TBD |
| K061EQ | Japan | Kagoshima | 29.8586°N | 129.833992°E | 10 | *Amphiprion frenatus* | TBD |
| K064 | Japan | Kagoshima | 31.5495°N | 130.647044°E | 6 | *Amphiprion clarkii* | TBD |
| K071 | Japan | Kagoshima | 31.549561°N | 130.647044°E | 10 | *Amphiprion clarkii* | TBD |
| K097 | Japan | Kagoshima | 30.464414°N | 130.499839°E | 12 | *Amphiprion clarkii* | TBD |
| K098 | Japan | Kagoshima | 30.462636°N | 130.500703°E | 19.7 | *Amphiprion clarkii* | TBD |
| K117 | Japan | Kagoshima | 30.814222°N | 131.029028°E | 21 | *Amphiprion clarkii* | TBD |
| K118 | Japan | Kagoshima | 30.814222°N | 131.029028°E | 21 | *Amphiprion clarkii* | TBD |
| K119 | Japan | Kagoshima | 30.814222°N | 131.029028°E | 21 | *Amphiprion clarkii* | TBD |
| K207 | Japan | Kagoshima | 31.62441111°N | 130.68323333°E | 11 | *Amphiprion clarkii* | TBD |
| K034EQ | Japan | Kagoshima | 28.06083°N | 129.27861°E | 2.6 | *Amphiprion frenatus* | TBD |
| K072 | Japan | Kanagawa | 35.294189°N | 139.556109°E | 2 | None | TBD |
| K074 | Japan | Kanagawa | 35.294189°N | 139.556109°E | 2 | None | TBD |
| K155 | Japan | Ogasawara Islands | 27.10122222°N | 142.23552777°E | 18.1 | *Amphiprion clarkii* | TBD |
| K156 | Japan | Ogasawara Islands | 27.10122222°N | 142.23552777°E | 17.8 | *Amphiprion clarkii* | TBD |
| K158 | Japan | Ogasawara Islands | 27.12111111°N | 142.1805°E | 29 | *Amphiprion clarkii* | TBD |
| K160 | Japan | Ogasawara Islands | 27.12111111°N | 142.1805°E | 24.7 | *Amphiprion clarkii* | TBD |
| K162 | Japan | Ogasawara Islands | 27.12111111°N | 142.1805°E | 19.3 | *Amphiprion clarkii* | TBD |
| K163 | Japan | Ogasawara Islands | 27.12111111°N | 142.1805°E | 22 | *Amphiprion clarkii* | TBD |
| K167 | Japan | Ogasawara Islands | 27.08141666°N | 142.18738888°E | 11.1 | *Amphiprion clarkii* | TBD |
| K168 | Japan | Ogasawara Islands | 27.08141666°N | 142.18738888°E | 18.1 | *Amphiprion clarkii* | TBD |
| K169 | Japan | Ogasawara Islands | 27.12025°N | 142.16297222°E | 15.7 | *Amphiprion clarkii* | TBD |
| K170 | Japan | Ogasawara Islands | 27.12025°N | 142.16297222°E | 9.6 | *Amphiprion clarkii* | TBD |
| K171 | Japan | Ogasawara Islands | 27.12025°N | 142.16297222°E | 23.2 | *Amphiprion clarkii* | TBD |
| K173 | Japan | Ogasawara Islands | 27.12025°N | 142.16297222°E | 14.8 | *Amphiprion clarkii* | TBD |
| K176 | Japan | Ogasawara Islands | 27.12025°N | 142.16297222°E | 20 | *Amphiprion clarkii* | TBD |
| K177 | Japan | Ogasawara Islands | 27.04133333°N | 142.20261111°E | 16 | *Amphiprion clarkii* | TBD |
| K178 | Japan | Ogasawara Islands | 27.04133333°N | 142.20261111°E | 22 | *Amphiprion clarkii* | TBD |
| K179 | Japan | Ogasawara Islands | 27.04133333°N | 142.20261111°E | 6.8 | *Amphiprion clarkii* | TBD |
| K184 | Japan | Ogasawara Islands | 27.04133333°N | 142.20261111°E | 13.5 | *Amphiprion clarkii* | TBD |
| K185 | Japan | Ogasawara Islands | 27.04133333°N | 142.20261111°E | 9.8 | *Amphiprion clarkii* | TBD |
| K186 | Japan | Ogasawara Islands | 27.05425°N | 142.17630555°E | 18 | *Amphiprion clarkii* | TBD |
| K187 | Japan | Ogasawara Islands | 27.05425°N | 142.17630555°E | 10 | *Amphiprion clarkii* | TBD |
| K189 | Japan | Ogasawara Islands | 27.11866666°N | 142.2295°E | 27.2 | *Amphiprion clarkii* | TBD |
| K190 | Japan | Ogasawara Islands | 27.11866666°N | 142.2295°E | 19.1 | *Amphiprion clarkii* | TBD |
| K191 | Japan | Ogasawara Islands | 27.11866666°N | 142.2295°E | 15.5 | *Amphiprion clarkii* | TBD |
| K192 | Japan | Ogasawara Islands | 27.11866666°N | 142.2295°E | 25.3 | *Amphiprion clarkii* | TBD |
| K194 | Japan | Ogasawara Islands | 27.13094444°N | 142.22127777°E | 22 | *Amphiprion clarkii* | TBD |
| K195 | Japan | Ogasawara Islands | 27.13094444°N | 142.22127777°E | 25.2 | *Amphiprion clarkii* | TBD |
| K199 | Japan | Ogasawara Islands | 27.18336111°N | 142.22127777°E | 18.5 | *Amphiprion clarkii* | TBD |
| K200 | Japan | Ogasawara Islands | 27.18336111°N | 142.22127777°E | 14.1 | *Amphiprion clarkii* | TBD |
| K202 | Japan | Ogasawara Islands | 27.18336111°N | 142.22127777°E | 12.9 | *Amphiprion clarkii* | TBD |
| K203 | Japan | Ogasawara Islands | 27.18336111°N | 142.22127777°E | 17.5 | *Amphiprion clarkii* | TBD |
| K205 | Japan | Ogasawara Islands | 27.18336111°N | 142.22127777°E | 16.7 | *Amphiprion clarkii* | TBD |
| K001 | Japan | Okinawa | 26.324722°N | 127.744444°E | 8.1 | *Amphiprion clarkii* | TBD |
| K003 | Japan | Okinawa | 26.324722°N | 127.744444°E | 6.7 | *Amphiprion frenatus* | TBD |
| K005EQ | Japan | Okinawa | 26.441666°N | 127.712222°E | 17.5 | *Amphiprion clarkii* | TBD |
| K006EQ | Japan | Okinawa | 26.441666°N | 127.712222°E | 10.0 | *Amphiprion clarkii* | TBD |
| K009EQ | Japan | Okinawa | 26.441666°N | 127.712222°E | 12.0 | *Amphiprion clarkii* | TBD |
| K016 | Japan | Okinawa | 26.537777°N | 128.079722°E | 6 | *Amphiprion frenatus* | TBD |
| KO68EQ | Japan | Sakurajima | 31.589981°N | 130.591964°E | 3.3 | *Amphiprion clarkii* | TBD |
| K018EQ | Japan | Shizuoka | 35.02164°N | 138.86691°E | 1.0 | None | TBD |
| K107 | Japan | Shizuoka | 35.02165°N | 138.86691°E | 2.0 | None | TBD |
| K108 | Japan | Shizuoka | 35.02166°N | 138.86691°E | 3.0 | None | TBD |
| K109 | Japan | Shizuoka | 35.02167°N | 138.86691°E | 4.0 | None | TBD |
| K078 | Japan | Wakayama | 35.454167°N | 135.748611°E | 20 | *Amphiprion clarkii* | TBD |
| K079 | Japan | Wakayama | 35.454167°N | 135.748611°E | 20 | *Amphiprion clarkii* | TBD |
| K082 | Japan | Wakayama | 35.454167°N | 135.748611°E | 10 | *Amphiprion clarkii* | TBD |
| K084 | Japan | Wakayama | 35.454167°N | 135.748611°E | 10 | *Amphiprion clarkii* | TBD |
| K085 | Japan | Wakayama | 35.454167°N | 135.748611°E | 12 | *Amphiprion clarkii* | TBD |
| MAL14EQ2 | Maldives | Fares-Maathoda | 0.262295°N | 73.214765°E | 16 | *Amphiprion clarkii* | TBD |
| MAL14EQ3 | Maldives | Fares-Maathoda | 0.262295°N | 73.214765°E | 16 | *Amphiprion clarkii* | TBD |
| MAL21EQ5C | Maldives | Fares-Maathoda | 0.262295°N | 73.214765°E | 16.3 | *Amphiprion clarkii* | TBD |
| MAL21EQ6C | Maldives | Fares-Maathoda | 0.262295°N | 73.214765°E | 16.5 | *Amphiprion clarkii* | TBD |
| MAL25EQ10B | Maldives | Fares-Maathoda | 0.262295°N | 73.214765°E | 16.3 | *Amphiprion clarkii* | TBD |
| MAL29EQ19C | Maldives | Fares-Maathoda | 0.262295°N | 73.214765°E | 16.3 | *Amphiprion clarkii* | TBD |
| PHILEQ1 | Philippines | NA | NA | NA | NA | NA | TBD |
| PHILEQ3 | Philippines | NA | NA | NA | NA | NA | TBD |
| SPEQ10 | Singapore | NA | 1.214892°N | 103.786258°E | NA | *Amphiprion frenatus* | TBD |
| SPEQ13 | Singapore | NA | 1.214892°N | 103.786258°E | NA | *Amphiprion frenatus* | TBD |
| SPEQ14 | Singapore | NA | 1.214892°N | 103.786258°E | NA | *Amphiprion frenatus* | TBD |
| SPEQ15 | Singapore | NA | 1.214892°N | 103.786258°E | NA | *Amphiprion frenatus* | TBD |
| SPEQ22 | Singapore | NA | 1.228586°N | 103.73101°E | NA | *Amphiprion frenatus* | TBD |
| SPEQ29 | Singapore | NA | 1.228586°N | 103.73101°E | NA | *Amphiprion frenatus* | TBD |
| SPEQ35 | Singapore | NA | 1.228586°N | 103.73101°E | NA | *Amphiprion frenatus* | TBD |
| SPEQ8 | Singapore | NA | 1.212172°N | 103.835263°E | NA | *Amphiprion frenatus* | TBD |
| SPEQ9 | Singapore | NA | 1.214892°N | 103.786258°E | NA | *Amphiprion frenatus* | TBD |
| SPEQ001 | Singapore | NA | 1.223837°N | 103.747009°E | NA | *Amphiprion frenatus* | TBD |
| SPEQ004 | Singapore | NA | 1.223837°N | 103.747009°E | NA | *Amphiprion frenatus* | TBD |
| SPEQ007 | Singapore | NA | 1.212172°N | 103.835263°E | NA | *Amphiprion frenatus* | TBD |

**Table S3.** List of publicly available bait-capture sequence data for sea anemones that were included in phylogenomic divergence-time analyses for the bubble-tip sea anemone *Entacmaea quadricolor*. Table includes information on Superfamily, Genus, and Species designations based on the currently accepted taxonomy. Sample size per species and per locality is denoted by *N.* Locality indicates country of origin for each sample. Accession Numbers (Accession #) refer to unique accession number from GenBank for that sample.

| Superfamily | Genus | Species | *N* | Locality | Accession # |
| --- | --- | --- | --- | --- | --- |
| Actinernoidea | *Synhalcurias* | *elegans* | 1 | Japanese Archipelago | SAMN13244953 |
| Actinioidea | *Actinostella* | *californica* | 2 | Baja, Mexico | SAMN39969072  SAMN39969074 |
| Actinioidea | *Actinostella* | *bradleyi* | 1 | Baja, Mexico | SAMN39969075 |
| Actinioidea |  | *flosculifera* | 2 | Bocas del Toro, Panama | SAMN39969076 SAMN39969073 |
| Actinioidea |  | *flosculifera* | 1 | Mexico | SAMN13244890 |
| Actinioidea | *Actinioidea* | sp. | 1 | Panama | SAMN07774921 |
| Actinioidea | *Actinia* | *equina* | 2 | Spain | SAMN13244941  SAMN39969089 |
| Actinioidea |  | *equina* | 1 | Iceland | SAMN39969092 |
| Actinioidea |  | *equina* | 1 | Ireland | SAMN39969090 |
| Actinioidea | *Actinia* | *tenebrosa* | 1 | New Zealand | SAMN39969091 |
| Actinioidea | *Anthopleura* | sp. | 1 | Panama | SAMN13244886 |
| Actinioidea | *Anthostella* | *stephensoni* | 1 | South Africa | SAMN13244945 |
| Actinioidea | *Bolocera* | *kerguelensis* | 1 | South Orkneys, Antarctica | SAMN13244888 |
| Actinioidea | *Bunodactis* | *octoradiata* | 1 | Patagonia, Chile | SAMN13244894 |
| Actinioidea | *Bunodosoma* | *granuliferum* | 1 | Panama | SAMN13244947 |
| Actinioidea | *Entacmaea* | *quadricolor* | 1 | Southern Great Barrier Reef, Australia | SAMN39969116 |
| Actinioidea |  | *quadricolor* | 1 | Northern Territories,  Australia | SAMN39969120 |
| Actinioidea |  | *quadricolor* | 1 | Fares-Maathoda, Maldives | SAMN39969118 |
| Actinioidea |  | *quadricolor* | 1 | Japanese Archipelago | SAMN39969119 |
| Actinioidea |  | *quadricolor* | 1 | Tonga | SAMN39969117 |
| Actinioidea | *Epiactis* | *georgiana* | 1 | South Orkneys, Antarctica | SAMN13244887 |
| Actinioidea | *Isactinia* | sp. | 1 | New Zealand | SAMN39969093 |
| Actinioidea | *Isosicyonis* | *alba* | 1 | Antarctica | SAMN07774928 |
| Actinioidea | *Stephanthus* | *antarcticus* | 1 | Antarctica | SAMN13244952 |
| Actinioidea | *Heteractis* | *aurora* | 3 | Fares-Maathoda, Maldives | SAMN39969077  SAMN39969078  SAMN39969079 |
| Actinioidea | *Phymanthus* | *crucifer* | 1 | Panama | SAMN13244907 |
| Actinioidea | *Radianthus* | *crispa* | 1 | Thuwal, Saudi Arabia | SAMN39969088 |
| Actinioidea |  | *crispa* | 1 | Gulf of Oman,  United Arab Emirates | SAMN39969082 |
| Actinioidea |  | *crispa* | 1 | Moorea, French Polynesia | SAMN39969084 |
| Actinioidea |  | *crispa* | 1 | Southern Great Barrier Reef,  Australia | SAMN39969083 |
| Actinioidea |  | *crispa* | 1 | Palau | SAMN39969087 |
| Actinioidea | *Radianthus* | *doreensis* | 1 | Philippines | SAMN39969080 |
| Actinioidea |  | *doreensis* | 1 | Japanese Archipelago | SAMN39969081 |
| Actinioidea | *Radianthus* | *magnifica* | 1 | Moorea, French Polynesia | SAMN39969104 |
| Actinioidea |  | *magnifica* | 1 | Northern Great Barrier Reef, Australia | SAMN39969103 |
| Actinioidea |  | *magnifica* | 1 | Scattered Islands | SAMN39969105 |
| Actinioidea |  | *magnifica* | 1 | Thuwal, Saudi Arabia | SAMN39969108 |
| Actinioidea |  | *magnifica* | 2 | Fares-Maathoda, Maldives | SAMN39969106  SAMN39969107 |
| Actinioidea | *Radianthus* | *malu* | 2 | Tonga | SAMN39969085  SAMN39969086 |
| Actinioidea | *Stichodactyla* | *gigantea* | 1 | Northern Great Barrier Reef,  Australia | SAMN39969113 |
| Actinioidea | *Stichodactyla* | *haddoni* | 1 | Sri Lanka | SAMN39969112 |
| Actinioidea |  | *haddoni* | 3 | Japanese Archipelago | SAMN39969109  SAMN39969110  SAMN39969111 |
| Actinioidea | *Stichodactyla* | *helianthus* | 3 | Bocas del Toro, Panama | SAMN13244891  SAMN39969114  SAMN39969115 |
| Actinioidea | *Stichodactyla* | *mertensii* | 1 | Fares-Maathoda, Maldives | SAMN39969101 |
|  |  | *mertensii* | 1 | Phillipines | SAMN39969100 |
| Actinioidea |  | *mertensii* | 1 | Scattered Islands | SAMN39969102 |
| Actinioidea | *Stichodactyla* | *tapetum* | 1 | Northern Great Barrier Reef,  Australia | SAMN39969098 |
| Actinioidea |  | *tapetum* | 1 | Vietnam | SAMN39969099 |
| Actinioidea | *Cryptodendrum* | *adhaesivum* | 1 | Northern Great Barrier Reef, Australia | SAMN39969094 |
| Actinioidea |  | *adhaesivum* | 1 | Moorea, French Polynesia | SAMN39969095 |
| Actinioidea | *Thalassianthus* | *hemprichii* | 2 | Northern Great Barrier Reef, Australia | SAMN39969096  SAMN39969097 |
| Actinostoloidea | *Actinostola* | sp. | 1 | South Orkneys, Antarctica | SAMN13244889 |
| Actinostoloidea | *Stomphia* | *didemon* | 1 | Washington, USA | SAMN13244908 |
| Actinostoloidea | *Halcampella* | *fasciata* | 1 | South Orkneys, Antarctica | SAMN13244892 |
| Actinostoloidea | *Sicyonis* | sp. | 1 | Antarctica | SAMN07774926 |
| Edwardsioidea | *Edwardsia* | *timida* | 1 | Ireland | SAMN13244884 |
| Edwardsioidea | *Nematostella* | *vectensis* | 1 | Unknown | SAMN02953687 |
| Metridoidea | *Antholoba* | *achates* | 1 | Patagonia, Chile | SAMN13244944 |
| Metridioidea | *Exaiptasia* | *diaphana* | 1 | Unknown | SAMN03839803 |
| Metridioidea | *Laviactis* | *lucida* | 1 | Panama | SAMN13244950 |
| Metridioidea | *Alicia* | *sansibarensis* | 1 | South Africa | SAMN13244943 |
| Metridioidea | *Lebrunia* | *neglecta* | 1 | Bocas del Toro, Panama | SAMN07774927 |
| Metridioidea | *Sagartiogeton* | *awii* | 1 | Antarctica | SAMN13244946 |
| Metridioidea | *Bunodeopsis* | *globulifera* | 1 | Bocas del Toro, Panama | SAMN07774923 |
| Metridioidea | *Diadumene* | *leucolena* | 1 | Brazil | SAMN13244966 |
| Metridioidea | *Galatheanthemum* | *cf. profundale* | 1 | Japanese Archipelago | SAMN13244949 |
| Metridioidea | *Halcurias* | *pilatus* | 1 | Chile | SAMN07774925 |
| Metridioidea | *Actinauge* | *richardi* | 1 | Ireland | SAMN13244965 |
| Metridioidea | *Phelliactis* | sp. | 1 | Ireland | SAMN13244964 |
| Metridioidea | *Metridium* | *senile* | 1 | Maine, USA | SAMN13244909 |
| Metridioidea | *Sagartia* | *troglodytes* | 1 | Ireland | SAMN13244951 |

**Table S4.** Akaike Information Criterion results for demographic models simulated in dadi for the Entacmaea quadricolor species complex. Model code refers to the models depicted in Figure 4. ΔAIC represents the difference in AIC when compared to the top model. w_i_=Akaike modle weights. Models are listed according to their AIC rank and the highest ranked model is highlighted.

| **Model** | **log-likelihood** | **AIC** | **ΔAIC** | **Model**  **Likelihood** | ***w_i_*** | **Evidence**  **Ratio** |
| --- | --- | --- | --- | --- | --- | --- |
| Q | -6365.62 | 12747.24 | 0 | 1 | 1 | 1 |
| H | -6728.1 | 13470.2 | 722.96 | 1.02e-157 | 1.02e-157 | 9.74e+156 |
| D | -6818.33 | 13652.66 | 905.42 | 2.45e-197 | 2.45e-197 | 4.06e+196 |
| B | -6830.51 | 13681.02 | 933.78 | 1.70e-203 | 1.70e-203 | 5.85e+202 |
| O | -6833.24 | 13682.48 | 935.24 | 8.22e-204 | 8.22e-204 | 1.21e+203 |
| C | -6892.78 | 13803.56 | 1056.32 | 4.19e-230 | 4.19e-230 | 2.38e+229 |
| L | -7002.12 | 14018.24 | 1271 | 1.01e-276 | 1.01e-276 | 9.86e+275 |
| E | -7207.97 | 14429.94 | 1682.7 | 0 | 0 | Inf |
| J | -7273.14 | 14562.28 | 1815.04 | 0 | 0 | Inf |
| K | -7327.82 | 14669.64 | 1922.4 | 0 | 0 | Inf |
| M | -7376.64 | 14767.28 | 2020.04 | 0 | 0 | Inf |
| I | -7392.27 | 14800.54 | 2053.3 | 0 | 0 | Inf |
| P | -7802.8 | 15619.6 | 2872.36 | 0 | 0 | Inf |
| A | -8066.71 | 16145.42 | 3398.18 | 0 | 0 | Inf |
| N | -8878.37 | 17770.74 | 5023.5 | 0 | 0 | Inf |
| G | -9028.49 | 18072.98 | 5325.74 | 0 | 0 | Inf |
| F | -9414.95 | 18843.9 | 6096.66 | 0 | 0 | Inf |

**Supplemental Figures**

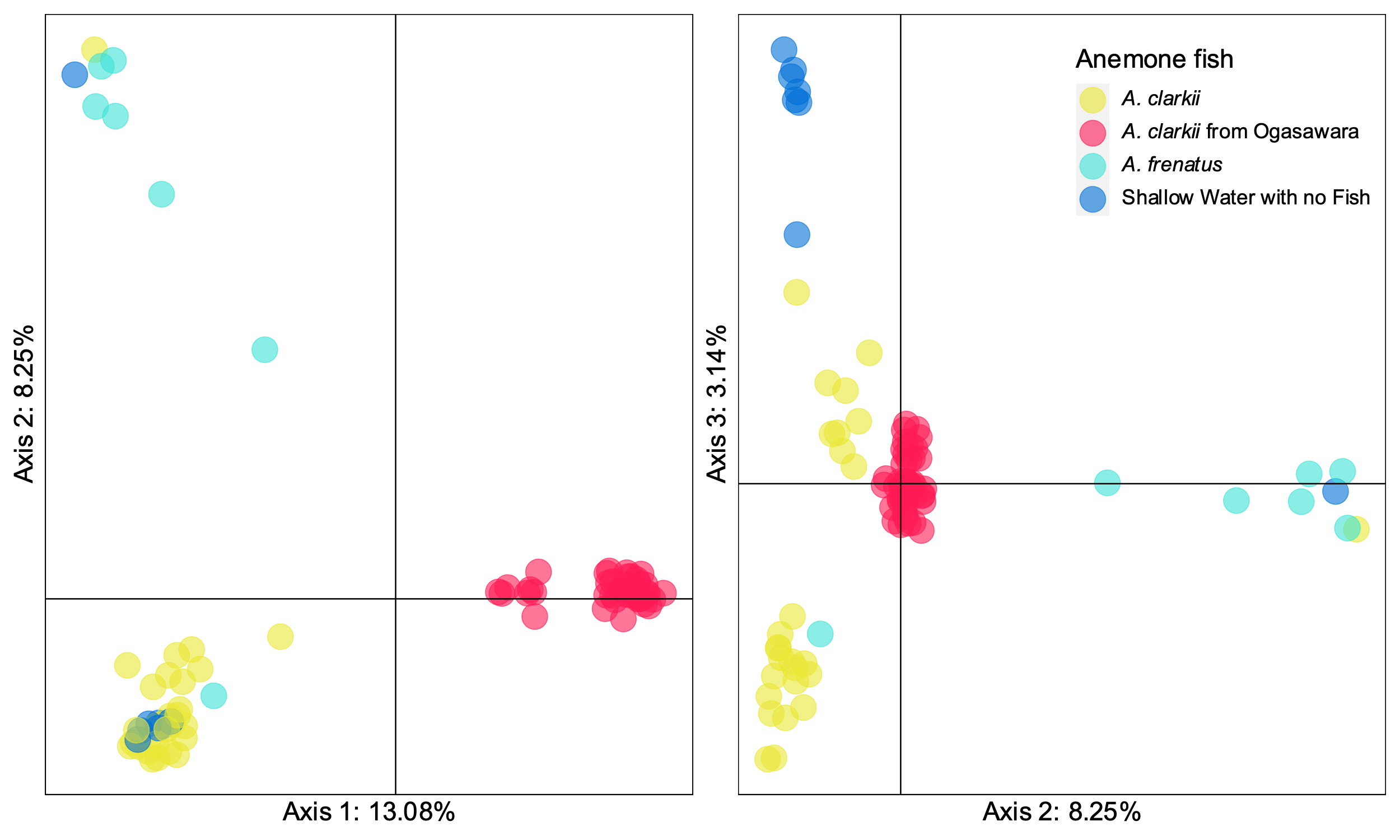

**Figure S1**. Principle Component Analysis (PCA) plots of the bubble tip sea anemone Entacmaea quadricolor from the Japanese Archipelago based on double digest Restriction site Associated DNA sequencing dataset. PCA reflects three major genetic partitions in the dataset, and is based on 3516 ddRADseq loci. Individual are colored by cryptic host lineage and the species of clownfish fish hosted by the individual. Anemones from Ogasawara (red) were colored differently from mainland Japan.

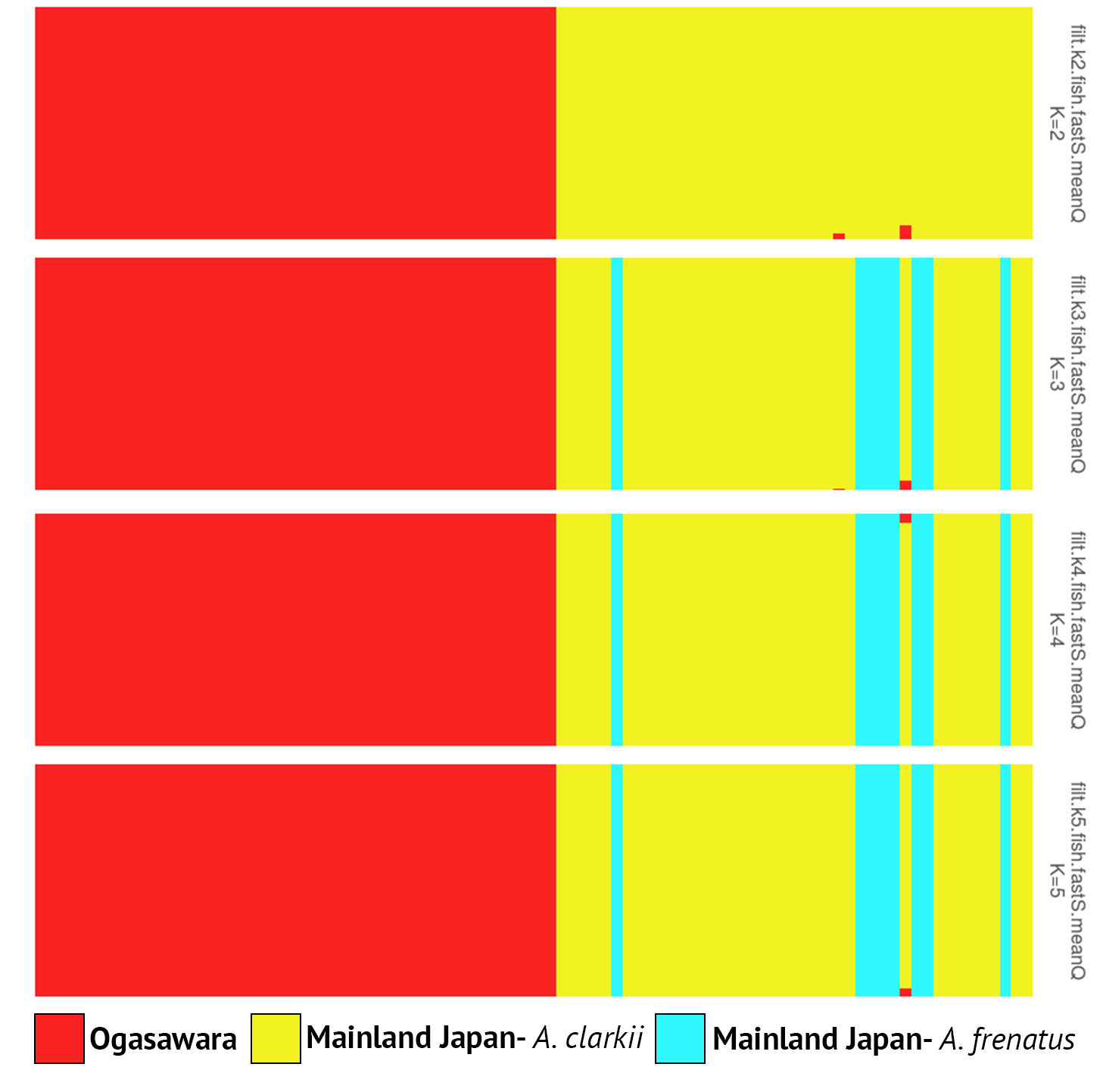

**Figure S2**. fastStructure genetic clustering results for *Entacmaea quadricolor* throughout the Japanese Archipelago using double digest Restriction site Associated DNA sequencing. Bar plot reflects *k*=2-5. Only three main genetic clusters were recovered in *k* = 3-5, which correspond to samples from Ogasawara Islands, and two co-occurring species in Mainland Japan that host *Amphiprion clarkii* and *A. frenatus* respectively.

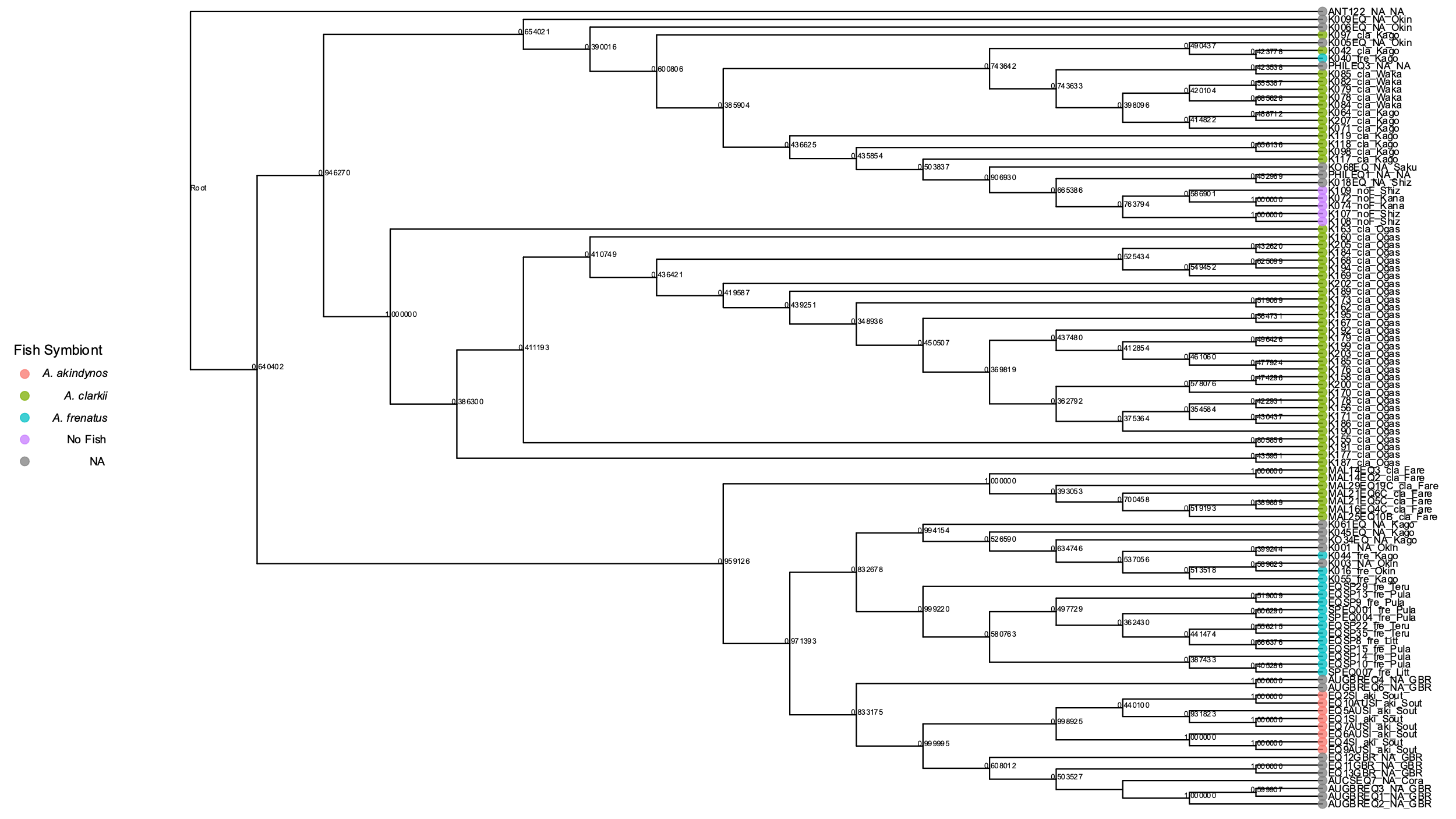

**Figure S3.** Coalescent-based phylogenetic reconstruction in ASTRAL-III of the bubble-tip sea anemone Entacmaea quadricolor. ASTRAL-III analyses using 1002 gene trees and 75% complete data matrix of ultra conserved elements and exon loci. Node support presented as posterior probabilities. Tip colors are based on fish symbiont species.

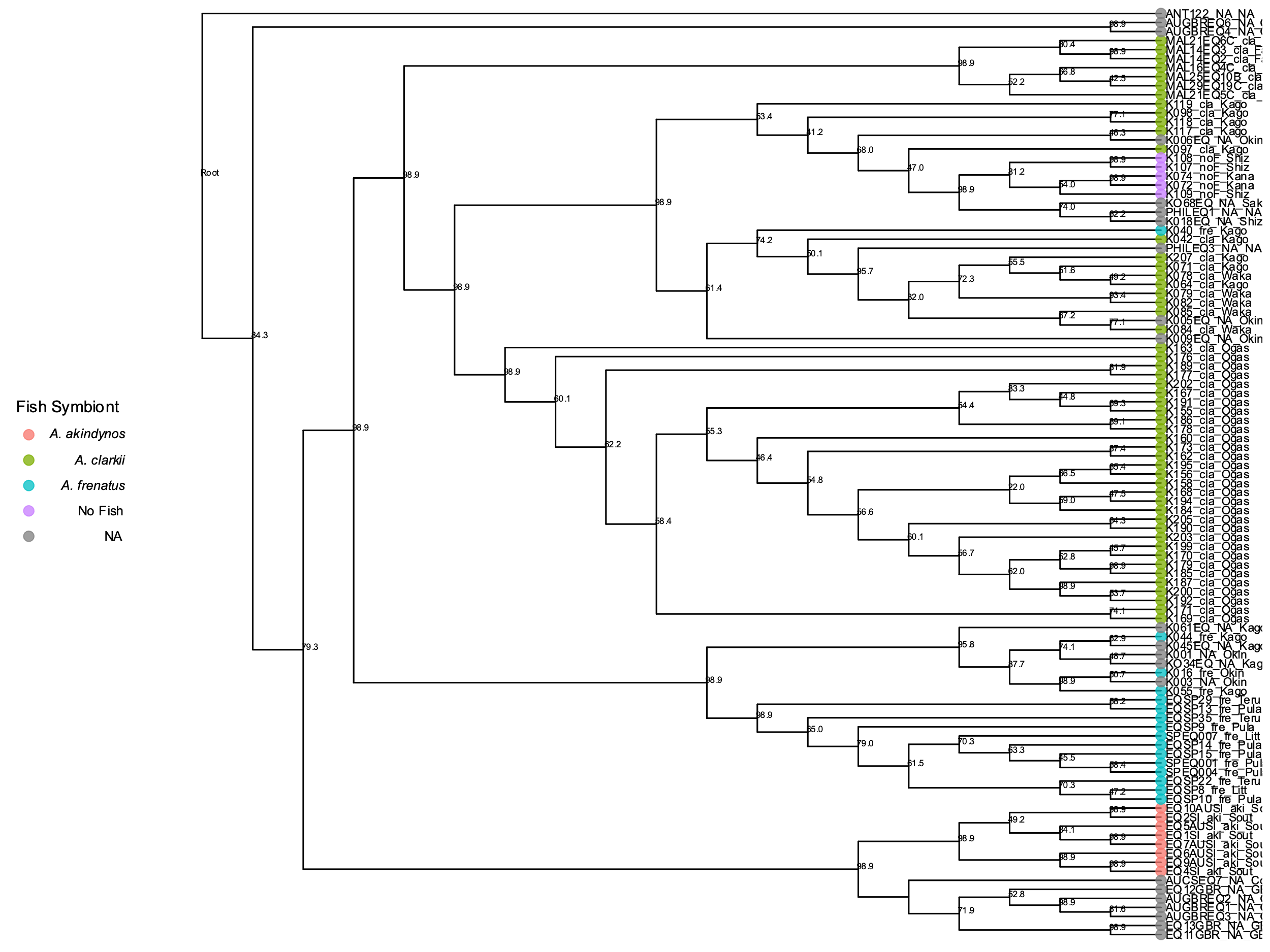

**Figure S4.** Coalescent-based phylogenetic reconstruction in CASTER of the bubble-tip sea anemone Entacmaea quadricolor. CASTER analyses using 1002 gene alignments and 75% complete data matrix of ultra conserved elements and exon loci. Node support presented as local bootstrap support. Tip colors are based on fish symbiont species

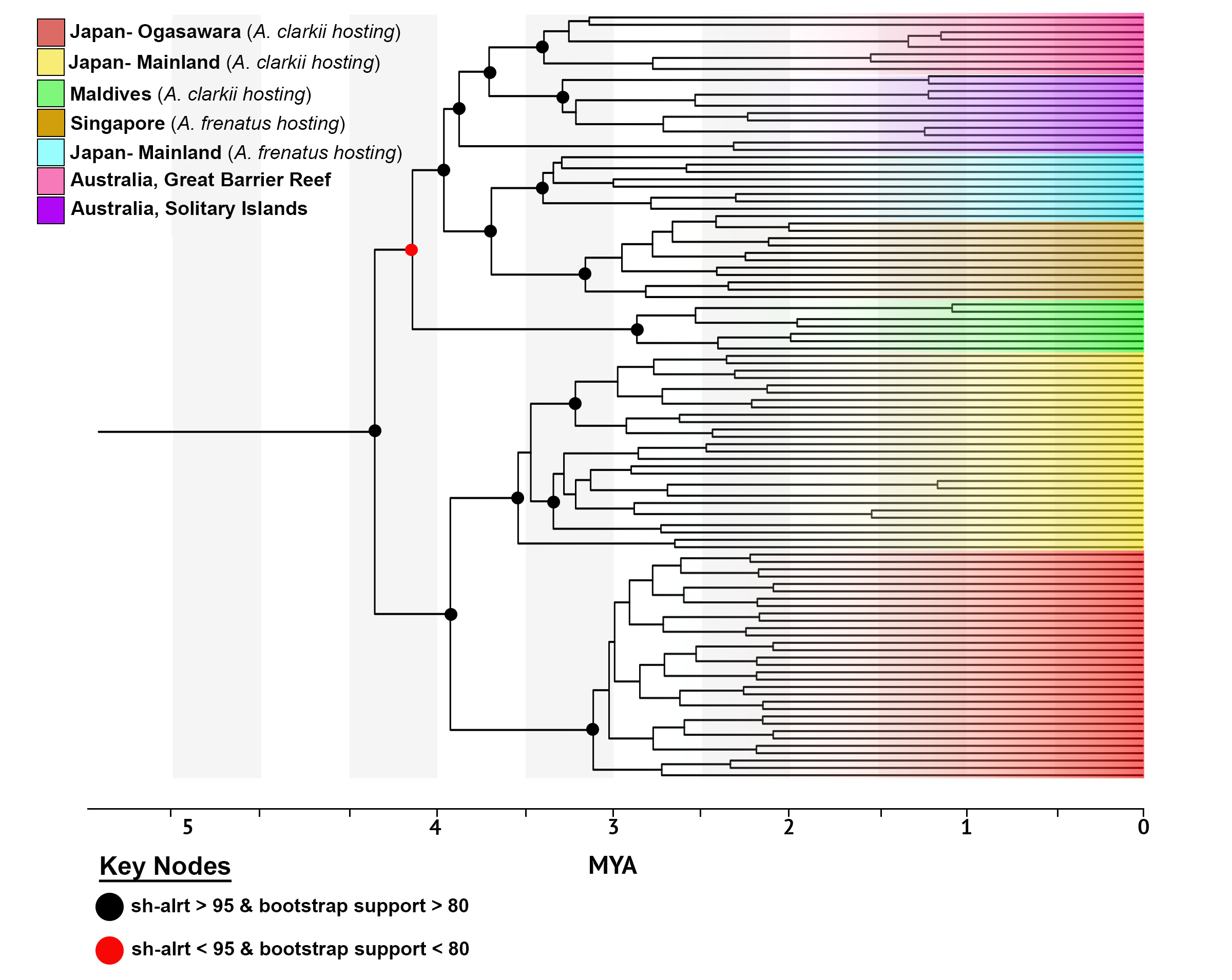

**Figure S5.** Time-calibrated maximum likelihood phylogenetic reconstruction for the bubble tip sea anemone *Entacmaea quadricolor*. ML analyses conducted using partitioned phylogenetic analysis in IQtree2 and bait-capture dataset targeting ultra-conserved element and exon loci (75% data occupancy matrix and 1002 loci). Scale bar represents millions of years before present (MYA).
